# The Regulatory-Associated Protein of the Target of Rapamycin Complex, RAPTOR1B, interconnects with the photoperiod pathway to promote flowering in *Arabidopsis*

**DOI:** 10.1101/2024.01.03.574092

**Authors:** Reynel Urrea-Castellanos, Maria J. Calderan-Rodrigues, Magdalena Musialak-Lange, Appanna Macharanda-Ganesh, Vanessa Wahl, Camila Caldana

**Author notes:** **Corresponding author:** Dr. Camila Caldana, Max-Planck-Institut für Molekulare Pflanzenphysiologie Am Mühlenberg 1; 14476, Potsdam-Golm (Germany). **Author Contributions:** R.U.C., V.W., and C.C. designed the research; R.U.C. performed the research; M.J.C.R. helped with selection of transgenic lines, inhibitor treatments and western blot analyses; M.M.L., and V.W. performed RNA in-situs; R.U.C. analyzed the data; R.U.C., and C.C wrote the paper. **Competing Interest Statement:** the authors declare no competing interest.

## Abstract

Transition from vegetative to reproductive growth (floral transition) is a strictly regulated energy-demanding process. In *Arabidopsis*, light perception coupled with internal circadian rhythms allows sensing changes in the duration of the light period (photoperiod) to accelerate flowering under long days (LD) in spring. This photoperiod-mediated floral induction relies on the accumulation of CONSTANS (CO) at dusk, a transcription factor that upregulates *FLOWERING LOCUS T* (*FT*) in leaves. Subsequently, FT protein moves into the shoot apical meristem to trigger the floral transition. Light and circadian clock-related signals are known to control CO at the genetic and protein levels; however, less is known about how energy sensing regulates components of the photoperiod pathway to modulate flowering. Here, we found that RAPTOR, a component of the Target Of Rapamycin complex (TORC), contributes to the induction of specific flowering genes that are under CO control. While transcription of *CO* remains intact in *raptor* mutants, its protein levels are reduced at dusk compared to wild-type (Col-0). This is due to increased protein degradation. Remarkably, GIGANTEA (GI) protein levels, which contributes to CO stabilization at dusk, are likewise hampered in the mutant. We show that RAPTOR interacts with and co-localizes at the nucleus with GI, altering GI levels through an unknown posttranscriptional mechanism. Phenotypic and molecular analysis of genetic crosses placed RAPTOR upstream of CO and GI. Since TORC is an energy sensor, our work suggests that RAPTOR could convey energy status information into the photoperiod sensing mechanism to fine-tune flowering behavior.

**Significance statement:** For annual plants, such as *Arabidopsis*, the correct timing of flowering in spring/summer is critical for reproductive success. Molecular mechanisms through which plants perceive and integrate day length with internal rhythms to accelerate flowering under long days are well described. However, little is known about the pathways sensing and conveying energy availability to the flowering programs. We found that RAPTOR, the regulatory unit of Target of Rapamycin complex (TORC), regulates CONSTANS post-transcriptionally through GIGANTEA. Both proteins are components of the photoperiod pathway of the flowering network and their miss-regulation in *raptor* mutants hampers the upregulation of genes promoting flowering. Our work suggests that the high energy available in long days is sensed and integrated into the photoperiod pathway by TORC.

## Introduction

Earth’s motion comprises rotation, which leads to a ∼24 h diel cycle of light and dark periods, and revolution, which changes the period’s length according to the season and latitude. For *Arabidopsis thaliana* (from now *Arabidopsis*) accessions that require vernalization (exposure to cold winter), sensing the photoperiod is crucial to maintain healthy vegetative growth in winter (1) and accelerate flowering during long days (LD) in spring (2). This timing of plant developmental transitions significantly impacts the production of offspring, aiming to grow and reproduce when the environmental conditions are favorable. Photoperiod-mediated flowering relies on the expression of the transcription factor CONSTANS (CO) in leaves, which in turn upregulates *FLOWERING LOCUS* (*FT*) (3, 4). FT is the florigen signal that moves from the leaves to the shoot apical meristem (SAM) to initiate floral transition (5). Accumulation of CO under LD results from an “external coincidence” mechanism that involves the circadian clock and photoperiod sensing (6). In short days (SD), clock-regulated *CO* gene expression and protein synthesis occur mainly after dusk, hindering CO accumulation since this protein is degraded in dark (6, 7). Contrary, in LD, high gene expression of *CO* coincides with the light period, where blue and far-red light photoreceptors stabilize further CO protein at dusk (7, 8). Additionally, the circadian-associated protein GIGANTEA (GI) interacts with FLAVIN-BINDING KELCH REPEAT F-BOX 1 (FKF1) and the circadian photoreceptor ZEITLUPE (ZTL) to regulate the diel protein turnover of CO in a time- and cell-specific manner (9). Thus, the FT-CO-GI module integrates circadian rhythms with light signals to accelerate flowering in LD(4).

Photosynthetic carbon (C) assimilation in the form of sugars (e.g., sucrose) occurs during the light period in plants. These sugars can be stored as starch or used as a readily available source of carbon (C availability) for growth (10). Long-term developmental changes, such as flowering, are predicted to be sensitive to C availability (11). Indeed, disruption of the sugar sensing mechanism mediated by trehalose 6-phosphate hampers flowering in *Arabidopsis*(12). In plants, major hubs integrate C availability information with external stimuli to control growth (13). The TARGET OF RAPAMYCIN (TOR) pathway has emerged as one of these hubs in different organisms, including plants (14). [NO_PRINTED_FORM]TOR is a protein kinase that associates with REGULATORY ASSOCIATED PROTEIN of TOR (RAPTOR) and LETHAL with SEC13 PROTEIN 8 (LST8) to form TOR complex 1 (TORC1) (15). A second complex (TORC2) exists in animals and yeast, where LST8 and additional proteins, with no homologs in plants, interact with TOR (16). In plants, the kinase activity of TOR leads to the activation of various connected metabolic processes that steer growth (17). In *Arabidopsis*, chemical inhibition of TOR activity or downregulation of *TOR* expression restricts overall growth (18–22), while knocking out TOR completely, or its kinase domain, leads to embryo lethality (23, 24). Although mutants of LST8 and RAPTOR display various developmental defects, they are viable and have been used to investigate the role of TORC (25–27). RAPTOR protein is encoded by two homologs, *RAPTOR1A* and *RAPTOR1B*; however, only knockout mutants for the later gene exhibit a delay in developmental transitions, including late flowering (28, 29). Remarkably, RAPTOR functions as the substrate recruiter for TOR, and its post-translational regulation through phosphorylation mediates the regulation of TOR kinase activity (30, 31), indicating that RAPTOR can integrate specific signals to selectively mediate TOR kinase substrates and/or its activity.

Since TORC malfunction has pleiotropic effects at the seedling stage (17), it is not clear if the late flowering associated with TORC inhibition, or with mutations in RAPTOR and LST8 genes (25, 27, 28), is only a consequence of early delays in development. Thus, to assess the role of TORC strictly at flowering, we took advantage of *RAPTOR1B* mutants given the fact that their growth behavior is very similar to wild-type under SD until the acquisition of floral competence. We found that RAPTOR promotes flowering under LD by regulating components of the photoperiod pathway. RAPTOR1B interacts with and promotes GI accumulation, thus contributing to CO stability at dusk, which in turn, mediates the upregulation of *FT* and other downstream genes required for floral transition. Favorable nutritional and C supply conditions activate TORC to rewire the metabolism and promote growth (32). Thus, growth defects upon disruption of TORC, such as in *raptor1b* and *lst8* cases, are more pronounced under LD (25)conditions (25) despite sufficient C availability (10). Therefore, our results suggest that plants incorporate C availability (energy status) information into the photoperiod pathway to properly regulate flowering under LD.

## Results

### RAPTOR positively regulates flowering

To understand the impact of the *RAPTOR1B* mutation during flower transition under LD conditions, two T-DNA insertion lines, named *raptor1b-1* and *raptor1b-2* (*SI Appendix*, Fig. S1A), were investigated. As previously reported (28, 29), both mutants bolted around 11 days later than wild-type (Col-0). However, we also observed an average of 8 extra leaves in the mutants under our growth conditions (Fig. 1A and B). In contrast, the knock-out mutant for *RAPTOR1A, raptor1a-1*, did not display any visible growth defects in our conditions (SI Appendix, Fig. S1B, C and D) or previous studies (28). Transformation of *raptor1b-1* plants with the *ProRAPTOR1B::6XMyc-RAPTOR1B or ProRAPTOR1B::RAPTOR1B-6XMyc* were able to fully restore the total leaf number and recovered the days of bolting from 87,8 % to 100% (*SI Appendix*, Fig. S2A and B).

**Figure 1.**
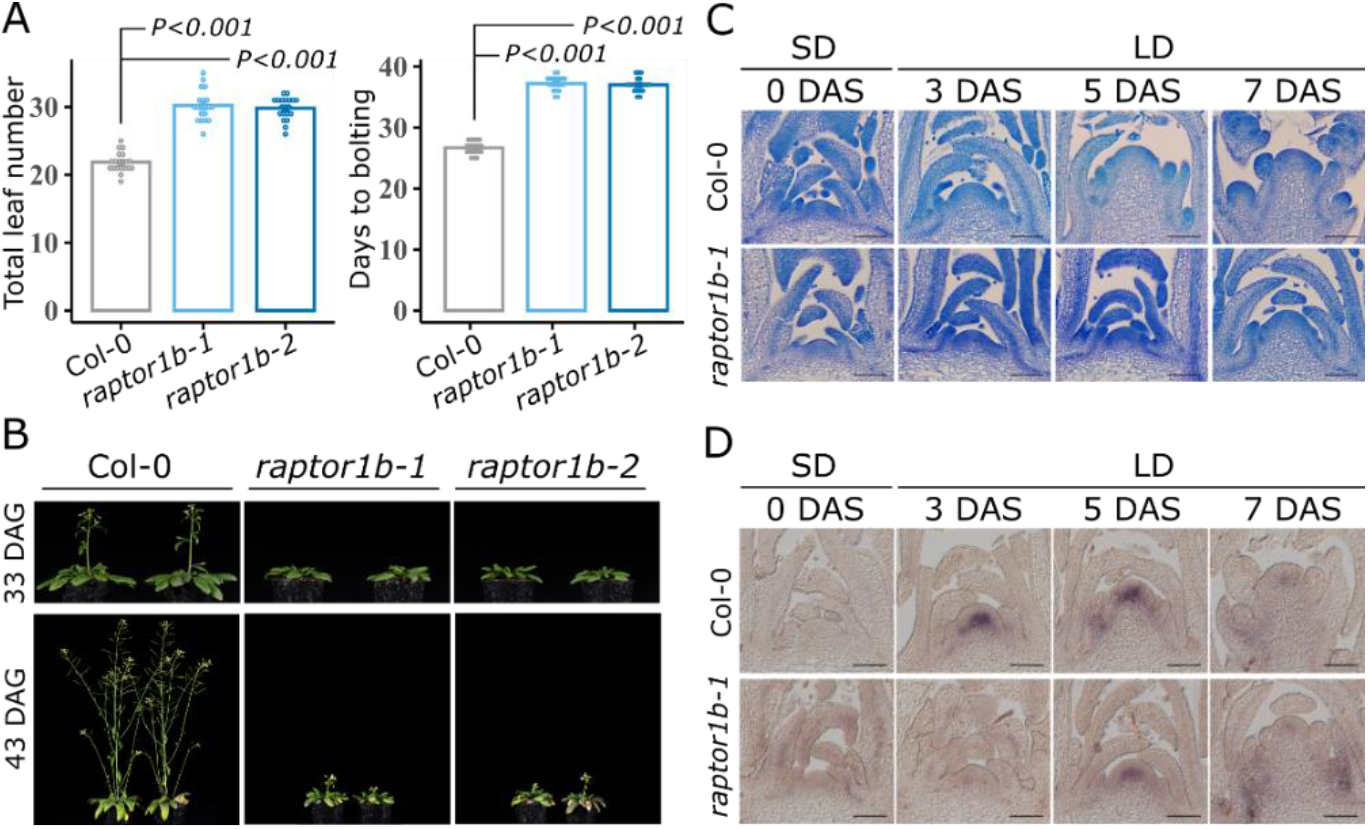
RAPTOR1B, the regulatory protein of the TOR complex, directly promotes flowering in *Arabidopsis* under LD conditions. **(A)** Days to bolting and total leaf numbers for Col-0 and two independent *raptor1b* mutant alleles, *raptor1b-1* and *raptor1b-2*, grown under LD. Significant differences between the genotypes were determined by two-tailed Student’s *t*-test (n= 20). Error bars denote Standard Error (SE). **(B)** Representative images of plants used to determine flowering time in (A). Pictures were taken 33 and 43 days after germination (DAG). **(C)** Longitudinal sections through Col-0 and *raptor1b-1* apices stained with Toluidine blue. Plants were first grown in SD for 30 days and then shifted to LD to induce flowering. Apices were harvested before (0 DAS) and after the shift (3, 5 and 7 DAS). Apices were collected 1h before dusk in SD and LD, respectively. DAS: days after shift. Scale bars: 100 μm. **(D)** RNA *in situ* hybridization using a specific probe for *SOC1* on Col-0 and *raptor1b-1* apices, for which plants were grown and harvested as in (C). DAS: days after shift. Scale bars: 100 μm.

Since *RAPTOR1B* mutants are affected early in development (26, 29), we investigated whether the late flowering phenotype could be a consequence of an overall developmental delay. To this aim, plants were grown first under SD for 30 days, when no phenotypic differences were observed between wild-type and *raptor1b-1* (*SI Appendix*, Fig. S3A), and then transferred to LD to induce flowering. [NO_PRINTED_FORM]Wild-type plants showed an increase in SAM size and formation of flower primordia at day 3 and 5, respectively, after the shift. Contrary, in *raptor1b-1*, SAM enlargement occurred much later, and no flower primordia were observed, thus confirming the late flowering phenotype of the mutant (Fig. 1C). The later induction and attenuated expression in the apices of *SUPPRESSOR OF OVEREXPRESSION CO1* (*SOC1*) in *raptor1b-1* compared to wild-type after the shift to LD indicated likewise a hampered floral transition in the mutant (Fig. 1D). Furthermore, the floral transition and *SOC1* expression at the SAM were delayed in continuous LD conditions even when the mutant seeds were sown two days in advance (staged *raptor1b-1*) to compensate for its later germination (*SI Appendix*, Fig. S3B and C) (26). Altogether, these results suggest that RAPTOR1B plays a direct role in the floral transition and that the late flowering phenotype of *raptor1b-1* is not due to delays in earlier developmental transitions.

### RAPTOR1B promotes the expression of specific flowering genes

We next measured the expression of key flowering genes in leaves of wild-type and *raptor1b-1* plants subjected to a photoperiod shift from SD to LD. As expected, this transfer induced *FT* expression in both genotypes (Fig. 2A). In *raptor1b-1*, however, *FT* levels were generally reduced, particularly at 5 and 7 days after the shift. Similarly, the closest homolog of *FT, TWISTER OF FT* (*TSF*), was induced under LD in both genotypes, but its expression in the mutant was affected in all time points (*SI Appendix*, Fig. S4A). The expression of *SQUAMOSA-PROMOTER BINDING PROTEIN-LIKE* (*SPL*) genes, in particular *SPL3, SPL4 and SPL5*, known to be induced in the rosette leaves and shoot apex upon induction to flowering (33), were likewise hampered in *raptor1b-1* upon transfer to LD (Fig. 2A). In contrast, other SPLs, such as *SPL9* and *SPL15*, were only significantly downregulated at day 7 in *raptor1b-1* compared to wild-type (*SI Appendix*, Fig. S4A). Reduced expression at dusk in *raptor1b-1* of *FT* and *SPL3, SPL4*, and *SPL*5, but not of *SPL9* or *SPL15*, was further confirmed over three consecutive diel cycles during the floral transition in continuous LD (*SI Appendix*, Fig. S5A). In addition, expression of *TOR* did not seem to be impaired in the *raptor1b-1* mutant. This is also the case for the *RAPTOR1B* gene (*SI Appendix*, Fig. S4A). Taking together, RAPTOR1B is required to promote the gene expression of positive regulators of flowering, namely *FT* and *SPLs* (*3, 4* and *5*), under LD conditions.

**Figure 2.**
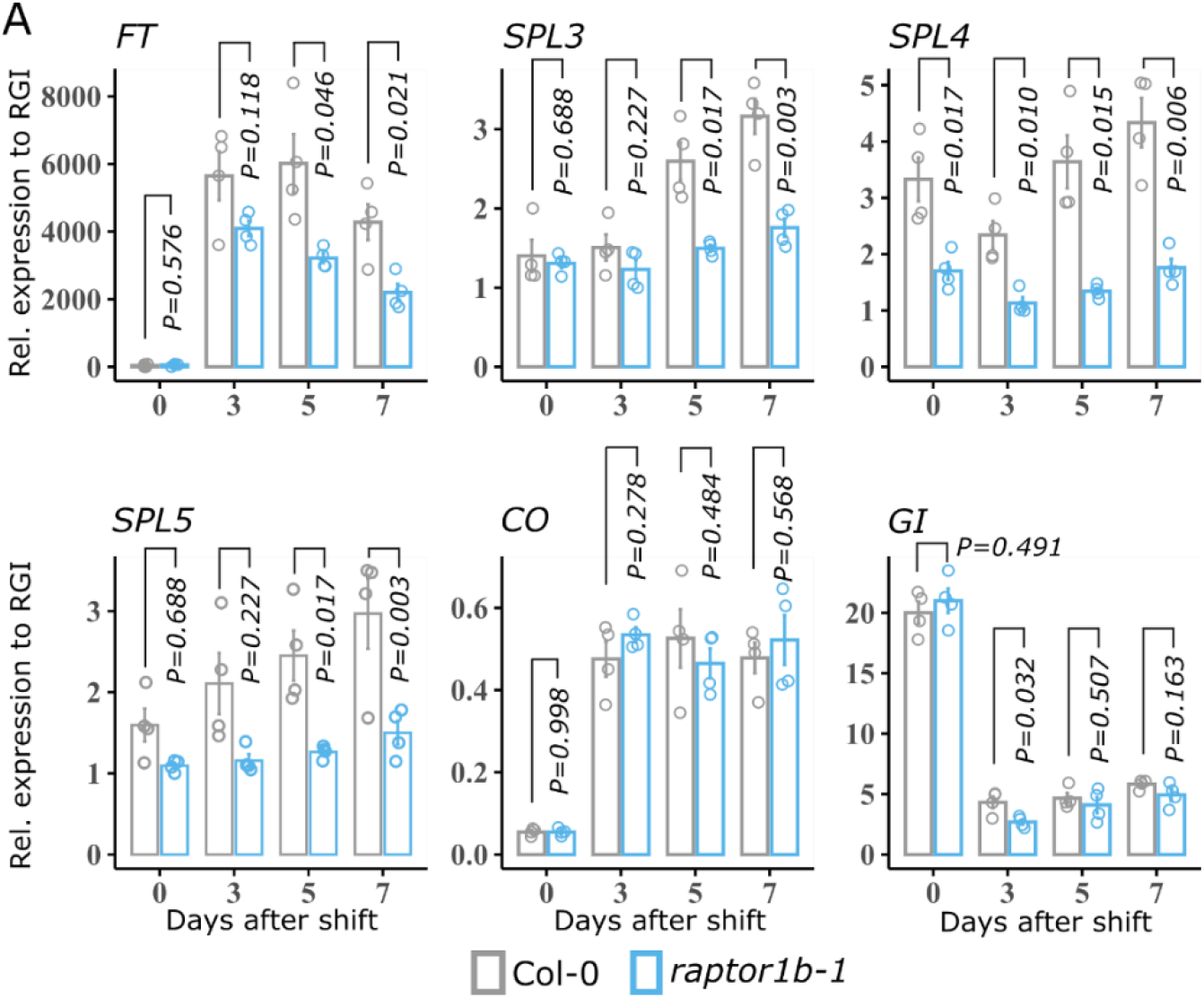
RAPTOR1B promotes the expression of flowering genes under LD conditions. **(A)** Gene expression analysis of *SPL3, SPL4, SPL5, FT, CO* and *GI* by RT-qPCR. Col-0 and *raptor1b-1* plants were grown initially in SD for 30 days and then transferred to LD. Rosettes were harvested before (0 DAS) and after the photoperiod shift (3, 5 and 7 DAS). Plant material was collected as described in Figure 1C. Significant differences between the genotypes for each day were determined by two-tailed Student’s *t-*test (n= 4). Error bars denote SE.

#### RAPTOR1B contributes to CONSTANS stability at dusk in long days

Expression levels of *FT, SPL3, SPL4, SPL5*, and *SOC1* are ultimately promoted by CO (8, 34). In contrast to the situation in SD, CO protein accumulates at dusk in LD as a result of both induced gene expression and light-mediated stabilization (6, 35, 36). *CO* expression itself was not affected in *raptor1b-1* neither at dusk throughout the SD to LD photoperiod shift (Fig. 2A) nor throughout a three-day time course under continuous LD (*SI Appendix*, Fig S5A). In contrast, *FT* expression was significantly downregulated in the mutant particularly at dusk in both experiments (Fig. 2A; *SI Appendix*, Fig. S5A). These results indicate that the reduced expression of flowering-promoting genes observed in *RAPTOR1B* mutant is not a consequence of *CO* gene expression.

Post-transcriptional regulation of the photoperiod pathway components plays a major role in regulating flowering. As expected, LD triggered the accumulation of CO in both genotypes. However, the protein abundance was reduced in *raptor1b-1* at all-time points upon shift to LD (Fig. 3A and B; *SI Appendix*, Fig. S6A), suggesting that RAPTOR1B might affect CO protein stability. To inspect the effect of the *raptor1b-1* mutation on CO protein regulation, we next applied the inhibitor of translation Cycloheximide (CHX) and the proteasome inhibitor MG132 to *raptor1b-1* and wild-type seedlings undergoing the floral transition using a hydroponic system[NO_PRINTED_FORM]Fig. 3C and D; *SI Appendix*, Fig. S6B). In wild-type, CO levels increased at dusk from day 9 to 10 after germination when the plants were grown in continuous LD, while for *raptor1b-1*, CO accumulation was hampered and reduced in comparison to wild-type. This observations resemble the differences in CO upregulation observed at the floral transition during the SD to LD shift experiment (Fig. 3A and B). CHX treatment did not significantly modify CO levels at dusk for both genotypes compared to the mock (DMSO), being CO levels in *raptor1b-1* still reduced when compared to wild-type. Contrary, under proteasome inhibition (+MG132), CO levels increased in *raptor1b-1* compared to the mock treatment and were more similar to the levels observed for wild-type (Fig. 3C and D; *SI Appendix*, Fig. S6B). The changes in CO accumulation under MG132 treatment suggest that CO protein degradation is enhanced in *raptor1b-1* at dusk. In summary, these results indicate that RAPTOR1B promotes floral transition under LD by facilitating the stabilization of CO at the end of the light period, and thus, accounts for the induction of flowering genes downstream of the photoperiod flowering pathway.

**Figure 3.**
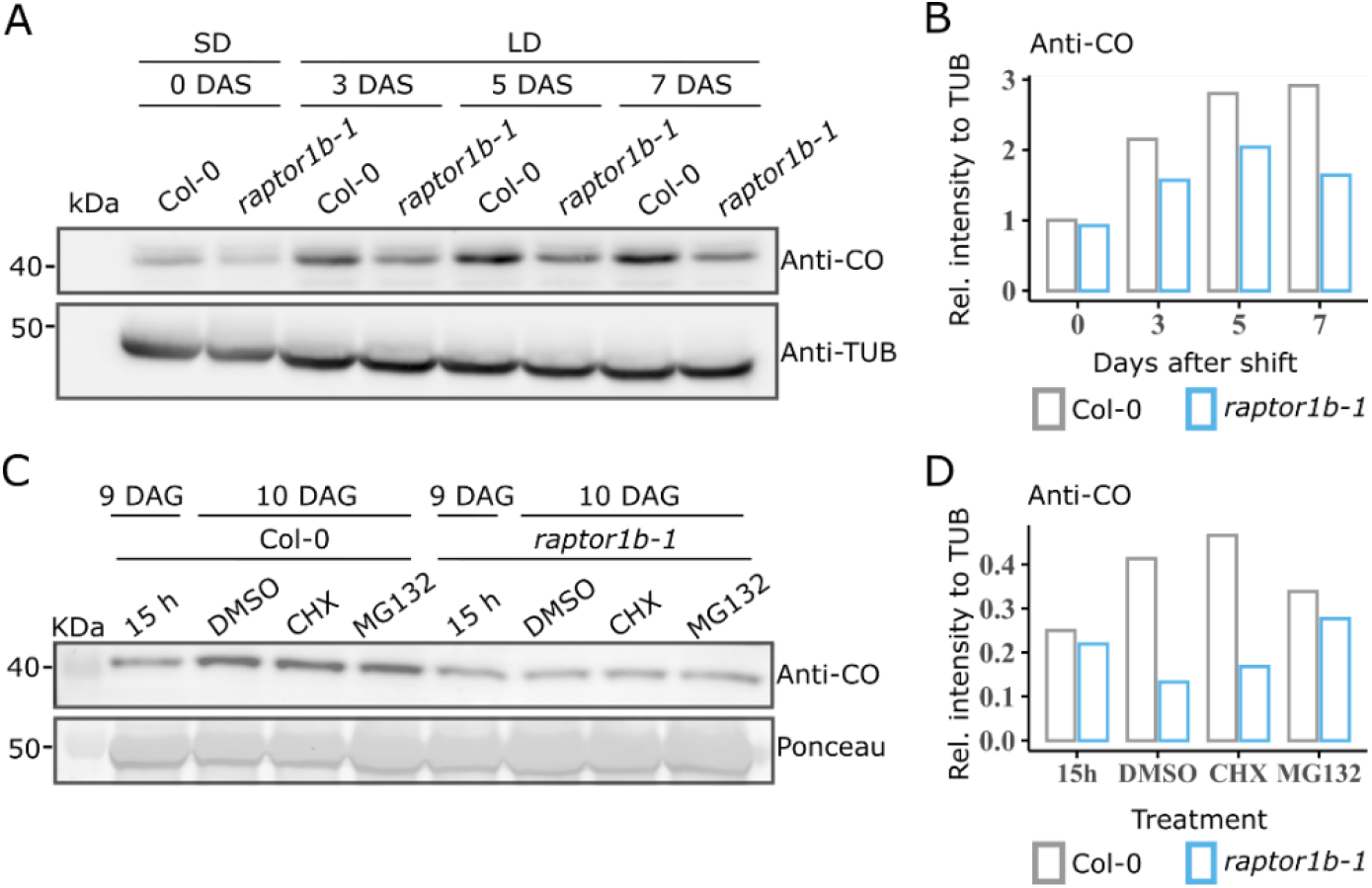
RAPTOR1B contributes to CONSTANS (CO) protein stability under LD. **(A)** Western blot analysis to determine protein levels of CO in Col-0 and *raptor1b-1* during a SD to LD shift experiment. Growth conditions and plant material are the same as described in Fig. 2A. Endogenous CO and TUBULIN proteins were immuno-detected using Anti-CO and Anti-TUB, respectively. One biological replicate is depicted here and the other three can be found in the *SI appendix*, Fig. S6A. **(B)** Relative CO protein levels calculated for the immunoblot in (A). Anti-CO was normalized to the respective Anti-TUB signal to determine the relative intensity at each day. **(C)** Western blot analysis of CO levels in Col-0 and *raptor1b-1* upon treatment with Cycloheximide (CHX, 100μM) or MG132 (100 μM). Plants were germinated and grown under LD in a hydroponic system for 10 days (at this time and conditions plants initiate the floral transition). Inhibitors and DMSO (1% v/v, mock treatment) were added 30 minutes before dawn. Entire rosettes were collected 15 h after the treatment at day 10. Shoots harvested at day 9 before the start of the treatment were used as control. Endogenous CO was immuno-detected with Anti-CO. A Ponceau stain was used as loading control. One biological replicate is depicted here, two more can be found in the SI Appendix, Fig. S6B. **(D)** Relative CO levels calculated for the immunoblot in (C). Anti-CO signal was normalized to the respective Ponceau loading control to determine the relative intensity for each treatment.

#### RAPTOR1B interacts with and accounts for GIGANTEA protein levels at dusk

We next investigated whether the altered CO stability in *raptor1b-1* was associated with light signaling (4). Blue light perception through CRYPTOCHROME 1 and 2 (CRY1 and CRY2) contributes to the CO abundance at dusk by suppressing CONSTITUTIVE PHOTOMORPHOGENIC 1 (COP1) and SUPPRESSOR of PHYA-105 1 (SPA1) mediated degradation (37–39). Likewise, far-red light perception via PHYTOCHROME A (PHYA) stabilizes CO (7), probably through the inactivation of the COP1-SPAs complex (4). PHYA and CRY1 protein levels were similar at dusk under SD and LD in both genotypes (*SI Appendix*, Fig. S7A and B), indicating no obvious connection between light sensing and the TORC pathway in regulating CO abundance. CO protein turnover is also regulated by a multilayered regulatory mechanism involving interaction among GI, FKF1, and ZTL (9). Given that GI acts by both stabilizing FKF1 and inhibiting ZTL to promote CO stability in the late afternoon, gene expression and protein levels of GI were assessed. *GI* transcription is not affected in *raptor1b-1b* (Fig. 2A and *SI Appendix*, Fig. S5A). Contrary, in both photoperiods, GI protein levels were reduced in the mutant compared to the wild-type (Fig. 4A and B; *SI Appendix*, Fig. S7C). Overall, these results suggest that RAPTOR1B might control the stability of CO via GI function at the post-transcriptional level.

**Figure 4.**
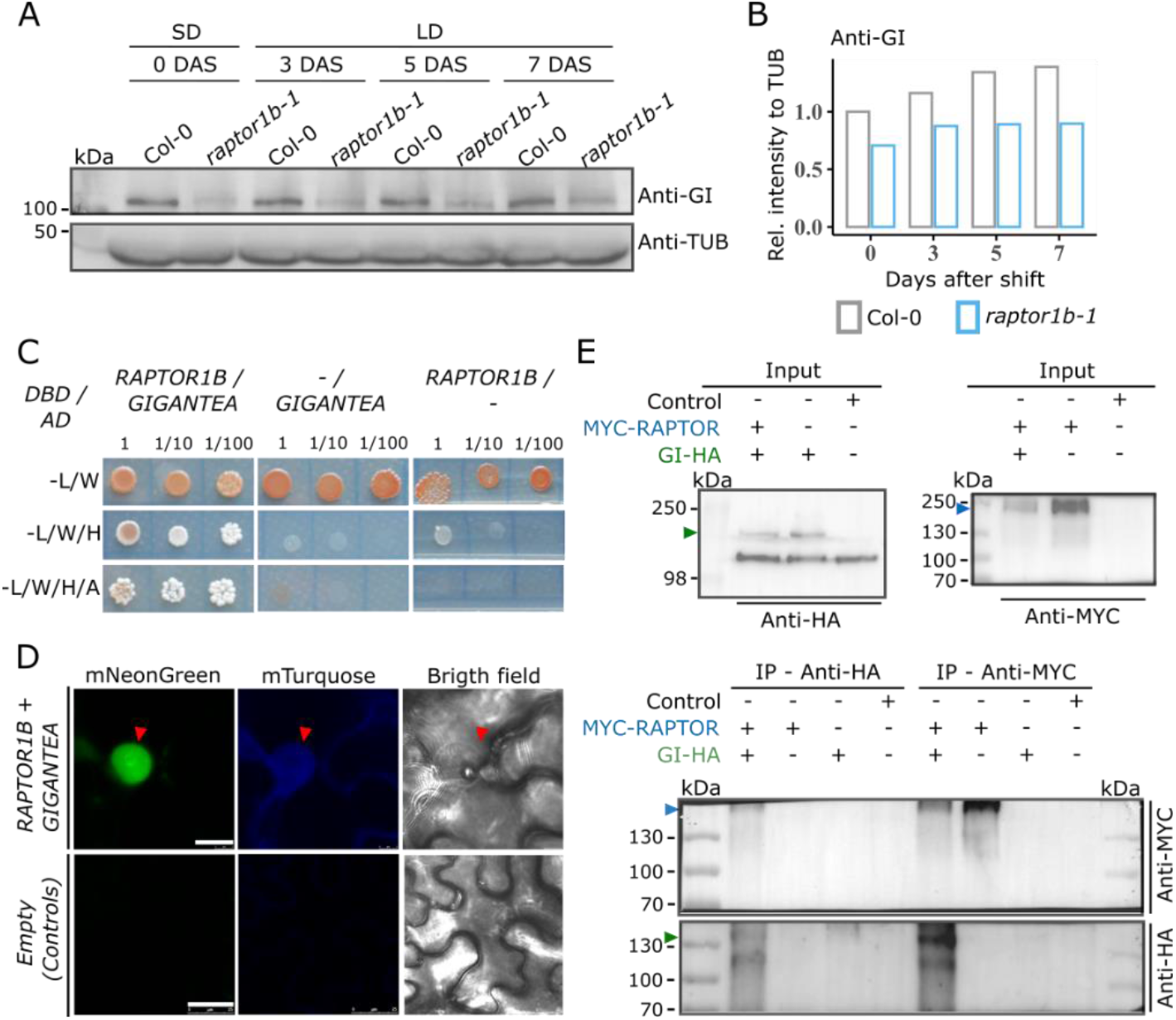
RAPTOR1B interacts with GIGANTEA (GI) and promotes its stability at dusk. **(A)** Western blot analysis to determine protein levels of GI in Col-0 and *raptor1b-1* grown in a SD to LD shift regime (as described in Fig. 2A). Endogenous GI and TUBULIN proteins were immuno-detected using Anti-GI and Anti-TUB, respectively. One biological replicate is depicted here, three more can be found in the *SI Appendix*, Fig. S7C. **(B)** Relative GI levels calculated for the immunoblot in (A). Anti-GI signal was normalized to the respective Anti-TUB signal to determine the relative intensity at each day. **(C)** RAPTOR1B and GI interact in yeast. Full sequences of *RAPTOR1B* and *GI* were used. Yeast was spotted on synthetic double (- L/W), triple (- L/W/H) or quadruple (- L/W/H/A) dropout medium. RAPTOR1B and S6K1 interaction was included as positive control (*SI Appendix*, Fig. S8A). L (Leucine), W (Tryptophan), H (Histidine), A (Adenine). **(D)** Transient co-expression of *Pro35S::mTurquoise-RAPTOR1B* and *Pro35S::GIGANTEA-mNeonGreen* in *N. benthamiana* leaves. As control, the empty vector (*pMDC32-HPB*) was infiltrated. Red arrow head indicates the nucleus. Scale bars, upper panel: 10 μm, bottom panel: 25 μm. Individual infiltration of both constructs was performed, respective images can be found in the *SI Appendix*, Fig. S9A and B. **(E)** Co-immunoprecipitation essays between RAPTOR1B and GI *in planta. Pro35S::6xMYC-RAPTOR1B* and *Pro35S::GI-3xHA* were transiently co-expressed in *N. benthamiana* leaves and protein extracts were immunoprecipitated (IP) with Anti-MYC or Anti-HA magnetic microbeads. Immunoblotting was performed using Anti-MYC and Anti-HA antibodies. As controls, *Pro35S::6xMYC-RAPTOR1B, Pro35S::GI-3xHA* and an unrelated protein (FLZ14-mNeongreen) were expressed individually (Control). Protein extracts prior to the immunoprecipitation (Inputs) are depicted in the top panels. Blue and green arrow heads indicate expected protein sizes for 6xMYC-RAPTOR1B (∼160 kDa) and GI-3xHA (∼132 kDa), respectively.

Interestingly, a yeast two-hybrid assay showed that RAPTOR1B interacted with GI and the 40S ribosomal protein S6 KINASE 1 (S6K1, positive control), but not with CO, FKF1 or ZTL (Fig. 4C; *SI Appendix*, Fig. S8A and B). Confirming the RAPTOR1B interaction with GI *in planta*, co-immunoprecipitation assays using *Pro35S:: 6XMYC-RAPTOR1B* and *Pro35S::GI-3XHA* showed that both recombinant proteins pulled-down each other when transiently co-expressed in *N. benthamina* leaves (Fig. 4E). To test for subcellular localization of this interaction, *Pro35S::mTurquose-RAPTOR1B* and *Pro35S::GI-mNeonGreen* were transiently co-expressed in *N. benthamina*. Both reporters were shown to co-localize in the nucleus (Fig. 4D). Contrary to GI-mNeonGreen, mTurquose-RAPTOR1B signal was also observed in the cytoplasm and cytoplasmic strings (*SI Appendix*, Fig. S9A and B). These results demonstrate that RAPTOR1B interconnects the TOR pathway with the photoperiod pathway through GI, contributing to CO stability at dusk.

#### GI and CO act downstream of RAPTOR1B

We next investigate the crosstalk between the RAPTOR1B and the photoperiodic pathway at genetic level. To this aim, we crossed *raptor1b-1* with previously reported lines expressing either *Pro35S::GI-TAP* or *ProCO::HA-CO*, aiming to increase the total amount of these proteins in the mutant background (see *SI Appendix for references*). Ectopic expression of HA-CO in *raptor1b-1* fully recovered the bolting time and total leaf number to Col-0 levels (Fig. 5A; *SI Appendix* Fig. S10A and S11A). However, these plants still flowered later when compared directly to the transgenic background used for the cross (*co-10,ProCO::HA-CO*), which shows the fastest bolting with the lowest total number of leaves of all the genotypes tested. On the other hand, ectopic expression of GI-TAP in *raptor1b-1* recovered partially both flowering time parameters but did not fully resembles the Col-0 phenotype. As observed for HA-CO, *raptor1b-1* expressing GI-TAP plants still flowered later in comparison to the transgenic background employed for the cross (*gi-2+Pro35S::GI-TAP*). Next, we tested if *raptor1b-1* does affect accumulation of GI-TAP and HA-CO in the crosses. Inspection of both chimeric proteins showed that their abundance was hampered likewise at bolting at dusk (*SI Appendix* Fig. S11A and B), as previously observed for the endogenous proteins (Fig 3A and Fig. 4A).

**Figure 5.**
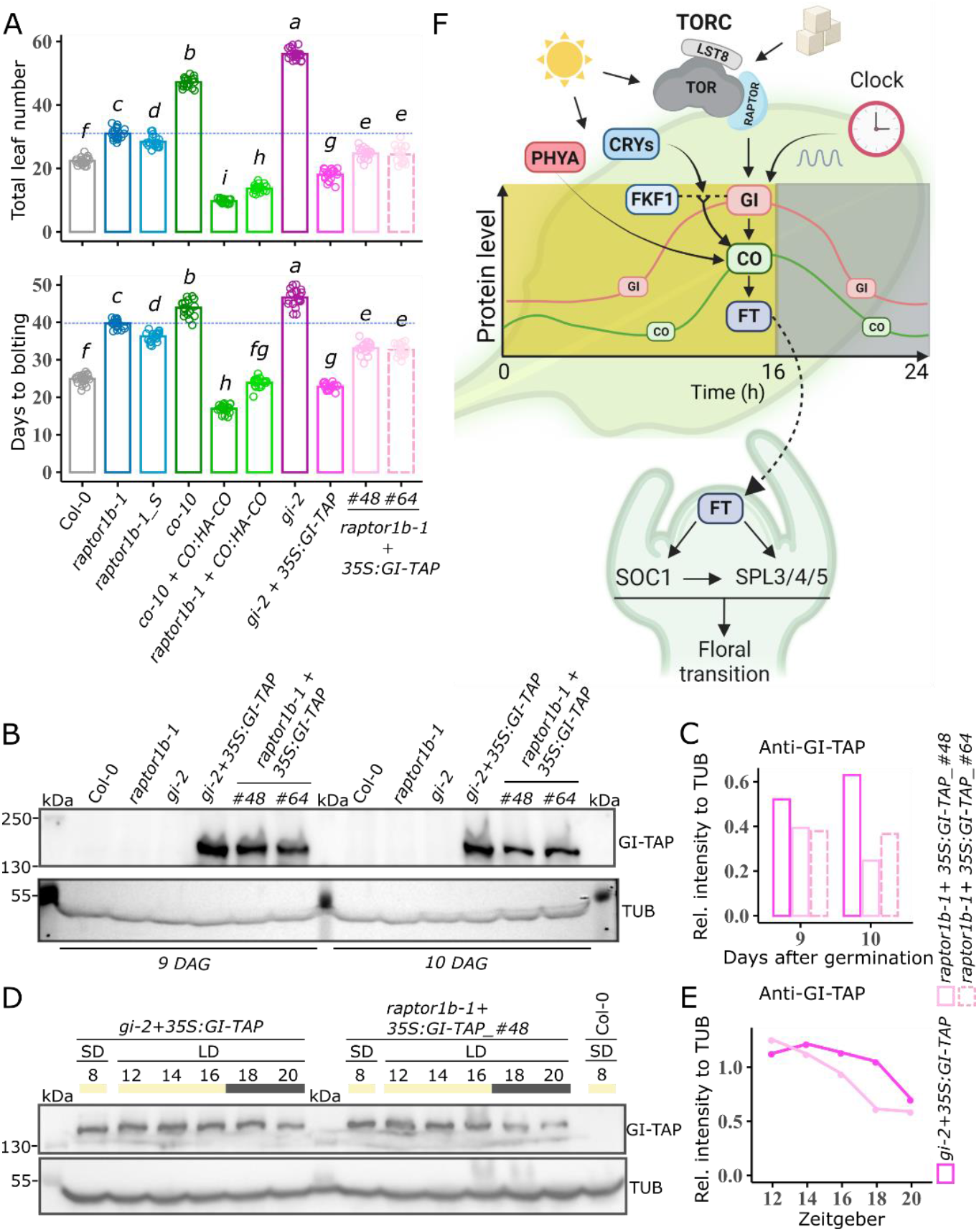
RAPTOR1B functions upstream of GI to contribute to CO stability at dusk and thus promote the floral transition in *Arabidopsis*. **(A)** Total leaf number and days to bolting under LD conditions for Col-0, *raptor1b-1, raptor1b-1_S* (staged), *co-10* (*constans* knock-out mutant), *co-10*+*ProCO:HA-CO* (complemented line for *co-10*), *raptor1b-1+ProCO:HA-CO* (*raptor1b-1* expressing HA-CO), *gi-2* (*gigantea* truncated mutant), *gi-2+Pro35S:GI-TAP* (complemented line for *gi-2*) and *raptor1b-1+Pro35S:GI-TAP* (*raptor1b-1* expressing GI-TAP, independent crosses: #48, #64). Significant differences among genotypes were determined by one-way ANOVA (*P* < 0.05) and a post-hoc Tukey’s test as indicated by letters (n=20). Error bars denote SE and a blue dotted line corresponds to the mean of *raptor1b-1*. **(B)** Western blot analysis of GI-TAP (∼150 kDa). Plants were grown in a hydroponic system. Whole rosettes were harvested in the last hour before dusk at 9 and 10 days after germination (DAG). GI-TAP was detected with Anti-GI and TUBULIN with Anti-TUB. One biological replicate is depicted here, three more can be found in the *SI Appendix*, Fig. S11C. **(C)** Relative GI-TAP levels calculated for the immunoblot in (B). Anti-GI signal was normalized to the respective Anti-TUB signal for relative intensity. **(D)** Western blot to determine GI-TAP abundance in *gi-2+35S:GI-TAP* and *raptor1b-1+Gi-TAP_#4*8 grown in a SD to LD shift regime. Rosettes were harvested at ZT8 in SD and every 2 h between ZT12 and ZT20 3 days after the shift in LD. Col-0 was included as control. Proteins were extracted and GI-TAP and TUBULIN were detected with Anti-GI and Anti-TUB, respectively. One biological replicate is depicted here and two more can be found in the *SI Appendix*, Fig. S12A. (**E**) Relative GI-TAP levels calculated for the immunoblot in (D). Anti-GI signal was normalized to the respective Anti-TUB for relative intensity. **(F)** Proposed model for the role of RAPTOR during floral transition in LD. The length of the light period is primarily sensed in leaves by the combined action of light signaling (mediated by PHYA and CRYs) and the circadian clock (driving gene expression of effector proteins, such GIGANTEA (GI) and FKF1). This accounts for protein accumulation of CONSTANS at dusk, which in turn, upregulates the florigen, *FT*. FT protein travels from the leaves into the shoot apex via the vasculature to induce the expression of floral integrator genes (such SOC1) and flower promoting genes (such SPL3/4/5). GI, similar to CO, accumulates over the day and is degraded at night. Independent, or together with FKF1 in response to blue light mediated by CRYPTOCHROMES (CRYs), GI helps to stabilize CONSTANS at dusk in LD. The Target of Rapamycin pathway, through RAPTOR, contributes to CO stability at dusk by promoting GI levels post-transcriptionally. The TOR complex (TORC) is known to be activated by light and sugars, suggesting that RAPTOR conveys energy status information to regulate GI accumulation and thus fine tune flowering via the photoperiod pathway.

Following, we investigated the impact of *raptor1B-1* mutation in the protein turnover of GI-TAP before and after dusk (from ZT12 to ZT20) upon a shift from SD to LD. In LD, *raptor1b-1* mutation hindered GI-TAP accumulation mainly at dusk and the first 4 h of the dark period when compared to ZT12 and ZT14 (Fig. 5D and E; *SI Appendix* Fig. S12A). These results indicate that in the presence of *raptor1b-1* mutation, GI-TAP decays faster in the late afternoon and the early period of the night. Confirming these observations, GI-TAP abundance at dusk was likewise dampened in the mutant background in plants undergoing the floral transition in a hydroponic system (Fig. 5B and C; *SI Appendix* Fig. S11C). Taken together, our results suggest that ectopic expression of GI-TAP or HA-CO compensates for the reduced levels of the endogenous proteins caused by the lack of RAPTOR. Likewise, but analysed from another perspective, introgression of *raptor1B-1* mutation into the background of the GI-TAP and HA-CO complementing lines diminishes their rescuing effect, indicating that GI and CO act downstream of RAPTOR.

## Discussion

We investigated the effect of the interplay between the cellular energy sensor TORC and the photoperiod pathway on the floral onset in *Arabidopsis*. A late flowering phenotype is associated with the malfunction of the TOR pathway (18), lack of RAPTOR (27, 28) or LST8 protein (25). However, since components of TORC also play a role in early developmental stages (17), it was first not clear if the TOR pathway directly impacts on flowering programs. That RAPTOR promotes the floral transition at the SAM independently of other pleiotropic functions at earlier developmental stages was suggested by a delayed conversion of the vegetative SAM into an inflorescence SAM, and attenuated expression of *SOC1* when *raptor1b-1* plants were subjected to a SD to LD shift (Fig. 1C and D), or staged to adjust development in LD (*SI Appendix*, Fig. S3B and C).

As a floral integrator, SOC1 perceives input from multiple flowering signals, such as age, vernalization, or photoperiod at the SAM (40). In [NO_PRINTED_FORM]response to LD, *SOC1* induction strongly reacts to the presence of FT, which integrates multiple flowering signals in leaves (3, 5). Generally, *SOC1* and *FT* respond strongest to signals reporting photoperiodic changes to adjust flowering time. In *raptor1b-1, FT* expression is still upregulated when exposed to inductive LD; however, transcript levels are significantly reduced at dusk compared to wild-type (Fig. 2A; *SI Appendix*, Fig. S5A). Similarly, mutants of *LST8*, the other accessory protein of the TORC, displayed hampered expression of *FT* after transfer from SD to LD conditions (25). Together with the bZIP transcription factor FD, FT directly, or via SOC1, induces *SPL3, SPL4*, and *SPL5* under LD (40, 41). This group of SPLs functions as co-activators of meristem identity genes (40, 42) and are upregulated in the rosette leaves and SAM upon induction to flowering (33). Hampered expression at dusk of *SPL3, SPL4*, and *SPL5* correlated with the downregulation of FT during the floral transition in the *raptor1b-1* mutant under LD (Fig. 2A; *SI Appendix*, Fig. S5A). [NO_PRINTED_FORM]

CO conveys multiple signals to induce the expression of *FT* and eventually *SOC*1 in LD and thus promote flowering (3, 4). Transcription of *CO* remained unaltered in *raptor1b-1* (Fig. 2A; *SI Appendix*, Fig. S5A). However, the consistent and significant downregulation of *SPL3, SPL4*, and *SPL5* at dusk, but not of other SPL genes, such as *SPL9* and *SPL15 (S*[NO_PRINTED_FORM]*I Appendix*, Fig. S4A and Fig. S5A), strongly pointed to a defect upstream of the photoperiod pathway (42)[NO_PRINTED_FORM]. In *raptor1b-1*, LD still induced CO accumulation at dusk, however, the protein levels were reduced when comparable to the wild-type (Fig. 3A and B; *SI Appendix*, Fig. S6A). To understand the post-transcriptional mechanisms responsible for the dampened CO levels in *raptor1b-1*, chemical inhibition of both translation and 26S proteasome-mediated degradation was performed. While inhibiting translation did not alter CO protein dynamics, blocking proteasome degradation brought CO closer to wild-type levels (Fig. 3C and D; *SI Appendix*, Fig. S6B), suggesting that RAPTOR accounts for CO stability at dusk. [NO_PRINTED_FORM][NO_PRINTED_FORM][NO_PRINTED_FORM]We investigated if the light-mediated stability of CO by CRY1/CRY2 and PHYA at dusk was affected in *raptor1b-1*; however no obvious differences in protein levels of the photoreceptors were observed compared to wild-type (*SI Appendix*, Fig. S7A and B). On the other hand, t[NO_PRINTED_FORM][NO_PRINTED_FORM]ogether with blue light perception through ZTL and FKF1 (43, 44), GI likewise accounts for CO accumulation at dusk (9). Transcript regulation of *GI* remained unaffected in *raptor1b-1* (Fig. 2A; *SI Appendix*, Fig. S5A); however, GI protein levels were reduced in the mutant (Fig. 4A and B; *SI Appendix*, Fig. S7C), suggesting a posttranscriptional miss regulation of GI. Interestingly, a delay in flowering time was achieved by enhanced degradation of GI in a temperature-dependent manner. This correlated with reduced *FT* expression, further indicating that posttranscriptional regulation of GI impacts flowering (45).

RAPTOR1B physically interacts with GI (Fig. 4C and E), but not with CO, FKF1 or ZTL (*SI Appendix*, Fig. S8A). *gi* mutants are late flowering in LD mainly due to downregulation of CO and FT (46), thus suggesting that RAPTOR1B promotes flowering via GI function under LD. Accordingly, ectopic expression of GI-TAP partially recovered the late flowering phenotype in the *raptor1b-1* background, while HA-CO expression, fully restored the flowering time parameters (Fig. 5A; *SI Appendix* Fig. S10A). Given that the complementing capacity of the GI-TAP transgene, as well as its protein levels, were hampered in presence of the *raptor1b-1* mutation (Fig. 5B, C, D and E), our results support the idea that RAPTOR1B control CO stability at dusk through GI function and places both photoperiod pathway components downstream of RAPTOR (Fig. 5F). CO stability depends on its phosphorylation status (47), and the TOR pathway can promote protein stability by regulating phosphorylation dynamics of downstream proteins, such in case of BETA BETA-AMYLASE 1 (BAM1) (48). However, it remains unclear if RAPTOR through GI could regulate CO accumulation through a similar process.

Also, the exact molecular mechanism by which RAPTOR controls GI stability remains unclear. A potential mechanism could involve C availability. In *raptor1b* mutants (29), or under TOR inhibition (19), sucrose levels in plants decreased, and this could result in lower GI levels since this sugar promotes GI accumulation (49). Thus, sucrose-dependent GI stability could be mediated by RAPTOR1B. Given that RAPTOR functions as the substrate recruiter for TOR kinase (14), GI could likewise be a phosphorylation target of the TOR pathway. Although in this study co-localization between RAPTOR and GI was only detected in the nucleus (Fig. 4D; *SI Appendix*, Fig. S9A and B), cytoplasmic localization of GI (44), RAPTOR (30) and TOR kinase (24) were previously reported. Indeed, GI-mediated flowering is regulated mainly by its nuclear localization (46, 50). Hence, the subcellular compartment of the interaction and the post-transcriptional modification of GI mediated by RAPTOR need to be investigated in the future. Remarkably, the TOR pathway and GI roles overlap for some biological processes, such as sugar-mediated entrainment of the circadian clock (51, 52). Furthermore, similar to mutants of the TOR complex (25, 27), mutations in *GI* lead to a starch excess phenotype in leaves (53). Consequently, our data support a scenario in which both the photoperiod and TOR pathways modulate plant growth, probably by mediating sugar signaling, via the GI-RAPTOR module.

## Material and Methods

### Plant material and growth conditions

All *Arabidopsis thaliana* plants used in this study are in the Columbia (Col-0) background. Reference of the mutants and description of the crosses used in this study are described in *SI Appendix*, Material and Methods. Detailed description of growth conditions for the flowering time and SD to LD photoperiod shift experiments, as well as for the chemical treatments with Cycloheximide (CHX, 100μM) and the proteasome inhibitor MG132 (100μM) performed in the hydroponics system, can be found in *SI Appendix*, Material and Methods.

### Generation of complementing transgenic lines in *Arabidopsis thaliana*

To complement the *raptor1b-1* mutant, 6XMyc was fused in frame either at the N-terminus or C-terminus of RAPTOR1B gene under the control of its endogenous promoter (*ProRAPTOR1B::6XMyc-RAPTOR1B, ProRAPTOR1B::RAPTOR1B-6XMyc*). Constructs were stably transformed by the floral dip method using *A. tumefaciens* strain GV3101. For details of plasmid construction and plant selection refer to *SI Appendix*, Material and Methods.

### Yeast two hybrid (Y2H)

Complete coding sequences of *RAPTOR1B* (*AT3G08850*), *GIGANTEA* (*AT1G22770*), *CONSTANS* (*AT5G15840*), *FKF1* (*AT1G68050*) and *ZTL* (*AT5G57360*) were used to generate the required Y2H constructs. For *S6K1* (*AT3G08730*), a region spanning from amino acid 9 to 293 was employed. Detailed information about plasmid construction and the conditions for the Y2H assay are found in *SI Appendix*, Material and Methods.

### Gene expression and RNA *in situ* analysis

For qRT-PCR, total RNA was extracted from approximately 50 mg of finely minced tissue in all cases using the Quick-RNA™ Plant Miniprep kit from ZYMO RESEARCH® (ref. R2024). Further details regarding cDNA synthesis, expression analysis and a list of primers used can be seen in *SI Appendix*, Material and Methods and Table S1. The probe to detect *SOC1* expression at the SAM as well as the *RNA in situ* hybridization method are described in *SI Appendix*, Material and Methods.

### Protein extraction and immunoblot analysis

50 mg of finely powdered tissue was suspended in 150 μl (3 volumes) of 2X extraction buffer (0.125 M Tris-HCl, pH 6.8; 4% SDS (v/v); 20% (v/v) glycerol; 0.01% (w/v) Bromophenol blue; 10% ß-mercaptoethanol), mixed vigorously and incubated at 95°C for 5 min. Denatured proteins were run in 8% (v/v) acrylamide gel containing 0.1% SDS, and then transferred into a PDVF membrane. Additional details about primary and secondary antibodies, running conditions, blotting, detection and analysis can be found in *SI Appendix*, Material and Methods.

### Co-Immunoprecipitation of proteins expressed in *Nicotiana benthamiana*

Full coding sequences of RAPTOR1B (*AT3G08850*) and GIGANTEA (*AT1G22770*) were fused with 6XMyc and 3XHA to generate *Pro35S::6XMyc-RAPTOR1B* and *Pro35S::GI-3XHA*, respectively. Constructs were transiently co-expressed in leaves of 6-week old *N. benthamiana* by leaf infiltration with *A. tumefaciens* strain GV3101. Proteins were extracted from 1g of finely minced tissue. Further details about protein extraction, co-immunoprecipitation and antibodies are found in *SI Appendix*, Material and Methods.

### Protein localization by transient expression in *Nicotiana benthamiana*

6-week old *N. benthamiana* leaves were infiltrated with *A. tumefaciens* strain GV3101 harboring *Pro35S::mTurquose-RAPTOR1B* or *Pro35S::GI-mNeonGreen*. Leaf pieces were visualized 48 h after infiltration using a SP5 confocal laser scanning microscope (©Leica Microsystems). Details about image acquisition and constructs are in *SI Appendix*, Material and Methods.

### Data visualization and statistical test

RStudio version 1.4.1717 was used for statistical analysis and plotting (https://www.rstudio.com). Statistical parameters are indicated in the legend of each figure. One-way ANOVA and Student’s *t-test* were performed by means of the R Stats package (https://stat.ethz.ch/R-manual/R-devel/library/stats/html/00Index.html). “Agricolae” package was used for the post-hoc Tukey’s test (https://cran.r-project.org/web/packages/agricolae/index.html). All plots were generated with help of the “ggplot2” package (https://cran.r-project.org/web/packages/DescTools/index.html).

## Supporting information

Supplemental Information Appendix of the main text

## Acknowledgments

We thank Patrick Giavalisco and Mohamed A. Salem for initial ideation and early support. We thank Alex Webb for providing seed material and Christin Abel for sectioning and plant care. Work in the Caldana lab was supported by the Max Planck Society. Work in the Wahl lab was funded by the BMBF (031B0191), the DFG (SPP1530: WA3639/1-2, 2-1), and the Max Planck Society.

