## Supplemental Information Appendix of the main text for "The Regulatory-Associated Protein of the Target of Rapamycin Complex, RAPTOR1B, interconnects with the photoperiod pathway to promote flowering in *Arabidopsis*"

***Corresponding author:**

Dr. Camila Caldana

**ORCID´s**

Reynel Urrea-Castellanos: https://orcid.org/0000-0002-0215-5423

Maria J. Calderan-Rodrigues: https://orcid.org/0000-0002-2422-8697

Magdalena Musialak-Lange: https://orcid.org/0000-0002-0388-8960

Appanna Macharanda-Ganesh

Vanessa Wahl: https://orcid.org/0000-0001-7421-8801

Camila Caldana: https://orcid.org/0000-0003-3639-0621


**This PDF file includes:**

Supplementary text
Figures S1 to S12
Table S1
SI References

**Supporting Information Text**

### Material and Methods

#### Plant material and growth conditions

All *Arabidopsis thaliana* plants used in this study are in the Columbia (Col-0) background. The mutant alleles for *RAPTOR1B* gene (*AT3G08850*), *raptor1b-1* (SALK_101990) and *raptor1b-2* (SALK_022096), were obtained from NASC and previously described (1, 2). *RAPTOR1A* gene (*AT5G01770*) mutant allele *raptor1a-1* (SALK_043920) was ordered from NASC and described before (1, 3, 4). *GIGANTEA* (*GI*) mutant *gi-2* and *GIGANTEA* complementing line *gi-2+Pro35S::GI-TAP* were provided by Prof. Alex Webb and described previously (5–8). *CONSTANS* (*CO*) mutant *co-10* (SAIL_24_H04) and complementing line *co-10+ProCO::HA-CO* were previously described (9, 10). *gi-2+Pro35S::GI-TAP* and *co-10+ProCO::HA-CO* were crossed with *raptor1b-1* (as pollen donor) and selected for homozygous *raptor1b-1* and presence of Pro*35S::GI-TAP* and *ProCO::HA-CO* constructs.

For flowering time experiments, seeds were first sown in a 1:1 mixture of soil (Stender) with vermiculite, stratified at 4°C for 3 days in darkness, and then transferred into a growth chamber. Plants were grown under a long photoperiod (LD, 16 h light / 8 h dark) with light intensity of 150 $\mu molm^{-2}s^{-1}$ and temperatures of 22°C / 18°C (light / dark). Bolting time was recorded when the inflorescence stem reached ≥ 0.5 cm. At this stage, total leaf number was determined after counting rosette and cauline leaves. For the short day (SD, 8 h light / 16 h dark) to long day (LD, 16 h light / 8 h dark) shift experiments, light intensity and temperature were kept at 160 $\mu molm^{-2}s^{-1}$ and at 22°C (day and night, to exclude the influence of temperature), respectively. Rosettes were harvested in the growth chambers and rapidly snap frozen in liquid nitrogen. Number of harvested individuals per replicate and time of the harvest are indicated in the figure legend of each experiment.

#### Hydroponic *in-vitro* experiment with MG132 and Cycloheximide (CHX)

For the *in vitro* experiment with the proteasome and translational inhibitors, Col-0 and *raptor1b-1* plants were grown for 10 days in a hydroponic system following the protocol described in (11). At this stage, plants are going throughout the floral transition. Growth conditions: LD (16 h light / 8 h dark) with temperatures of 21°C (day) and 19°C (night) and HR of 60%. 10 days after germination, 100µM cycloheximide (Sigma-Aldrich, cat no. C7698), 100µM MG132 (Sigma-Aldrich, cat no. 4747908) or DMSO (1 % v/v) (mock control) were added to the media 30 minutes before dawn. Next, shoots were harvested 15 h after dawn either before or after the treatment and snap-frozen in liquid nitrogen and kept at –80°C for further analysis.

#### Generation of complementing transgenic lines in *Arabidopsis thaliana*

PCR amplification was carried out using Phusion™ High-Fidelity DNA Polymerase (Thermo Scientific, ref. F-530XL). A list of primers is provided in Table S1. *ProRAPTOR1B::RAPTOR1B-6xMyc* construct was generated by amplifying first the coding sequence of *RAPTOR1B* ($RAPTOR1B_{cds}$, lacking the stop codon) from *A. thaliana* (Col-0) cDNA and cloned via *In-Fusion*® (Takara Bio, ref 638948) in frame into the *pE3c* vector before the *6XMyc* tag (Addgene, https://www.addgene.org). Next, the promoter region of *RAPTOR1B* (1363 bp fragment upstream from the start codon) was amplified from genomic DNA and cloned via *In-Fusion*® cloning upstream of $RAPTOR1B_{cds}$*-6xMyc* in the pE3c plasmid to generate *ProRAPTOR1B::RAPTOR1B-6xMy.* Subsequently, via Gateway cloning (Invitrogene™), *ProRAPTOR1B::RAPTOR1B-6xMy* was subcloned into the plant expression vector *pGWB501* (Addgene, https://www.addgene.org), which harbors the Hygromycin resistance gene for plant selection. For *ProRAPTOR1B::6xMyc-RAPTOR1B* construct, $RAPTOR1B_{cds}$ was amplified conserving the stop codon and cloned in frame into the *pE3n* vector downstream of the 6XMyc tag (Addgene, https://www.addgene.org). Following, cloning of the promoter region and subcloning into the plant destination vector, *pGWB501*, was done as described above. Both constructs were transformed into the *raptor1b-1* mutant background by the floral dip method (12) using *A. tumefaciens* (strain GV3101). Selection of positive transformants and homozygous transgenic lines in the F3 was performed in ½ MES media containing Hygromycin as described in (13). At least three independent transgenic lines for each construct were chosen for further analysis.

**Yeast two hybrid**

Full coding sequences of all genes were first cloned into the entry vector *pDONR221™* using Gateway cloning (Invitrogene™). Next, *RAPTOR1B* was recombined into *pDEST™32* (Invitrogene™), which harbors the DNA Binding Domain (DBD) at the N-terminal. *GIGANTEA*, *CONSTANS, FKF1, ZTL* and *S6K1* were recombined into *pDEST™22* (Invitrogene™), which harbors the Activation Domain (AD) at the N-terminal. The yeast strain Y2HGold (Takara Bio, ref. 630498) was double transformed with the bait plasmid (*DBD-RAPTOR1B*) in combination with each of the prey plasmids (*AD-GIGANTEA*, *AD-CONSTANS*, *AD-FKF1*, *AD-FKF1* and *AD-S6K1*) following the protocol in (14). As negative controls, empty *pDEST™32* and *pDEST™22* were likewise co-transformed with the prey and bait plasmids, respectively. Positive transformants were selected in double dropout media lacking Leucine and Tryptophan (- L/W) (Taka Bio, ref. 630495). Next, to test the interaction between *RAPTOR1B* and the prey proteins, recovered colonies from the previous selection were resuspended in 1X TE buffer and spotted into double (-L/W), triple (- L/W/H) and quadruple (- L/W/H/A) dropout media, following the protocol in (14). Primers are listed in Table S1.

**RNA extraction and gene expression analysis**

Total RNA was extracted from approximately 50 mg of finely minced tissue using the Quick-RNA™ Plant Miniprep kit from ZYMO RESEARCH® (ref. R2024). 1.8 -2 µg of total RNA was used for cDNA synthesis using the RevertAid H Minus First Strand cDNA Synthesis Kit from ThermoFisher (ref. K1632) and Oligo(dT)$_{18}$ as primer. 1 to 10 dilution of the synthesized cDNA was employed for further analyses. For gene expression analysis, qRT-PCR was carried out by means of Power SYBR Green PCR-Master_Mix (Applied Biosystems, ref. 4367659) and the ABI PRISM 7900HT system (Applied Biosystems) for detection. Selection of reference genes was performed for each type of experimental setup as described in (15), and the geometric mean of two selected reference genes (RGI) was used for further normalization. The comparative cycle threshold (CT) method was employed to determine the relative expression of the selected genes (16). In short, relative expression of the target gene was first normalized to the RGI to generate the ΔCt, and then normalized to the maximum value (referred as Max_Calibrator) to calculate the ΔΔCt. Next, final expression was computed following the $2^{-\Delta\Delta Ct}$ formula. Four biological replicates per time point and genotype were performed by harvesting a pool of ~5 plants in each case. Significant differences were determined by Student's *t-test*. Information of the primers can be found in Table S1.

**RNA *in situ* hybridization**

The probe to detect *SOC1* expression at the SAM as well as the *RNA in situ* hybridization method were previously described in (17, 18).

**Protein extraction and immunoblot analysis**

50 mg of finely powdered tissue was suspended in 150 µl (3 volumes) of 2X extraction buffer (0.125 M Tris-HCl, pH 6.8; 4% SDS (v/v); 20% (v/v) glycerol; 0.01% (w/v) Bromophenol blue; 10% ß-mercaptoethanol), mixed vigorously and incubated at 95°C for 5 min. Next, samples were centrifuged twice at 13,000 g for 5 min to remove cell debris. Denatured proteins were loaded and run into an 8% (v/v) Acrylamide gel containing 0.1% (v/v) SDS using a Bio-Rad Mini-PROTEAN Tetra System. Next, separated proteins were transferred into a 0.45 µm PDVF Immobilon-P membrane (Merck Millipore, ref. IPVH00010) and the PDVF membrane was blocked by incubation for 2 h at RT with 1X TBS-T buffer (20 mM Tris, 150 mM NaCl, pH 7.6, 1 ml.L$^{-1}$ Tween20) supplemented with 5% fat free milk. Primary antibodies, Anti-CONSTANS (PhytoAB, ref. PHY2297; 1:1000), Anti-GIGANTEA (Agrisera, ref. AS121864A; 1:1000), Anti-TUBULIN (Sigma-Aldrich, ref. T5168; 1:10000), Anti-CRY1 (PhytoAB, ref. PHY1707S; 1:1000) and anti-PHYA (PhytoAB, ref. 1907; 1:1000), were added and incubated with the membrane in 1X TBS-T buffer supplemented with 1% fat free milk overnight at 4°C. Following, after washing out the primary antibody 3 times with 1X TBS-T buffer, PDVF membrane was incubated with the secondary antibody, either Goat Anti-Rabbit (Bio-Rad, ref. 1706515) or Anti-Mouse (Bio-Rad, ref. 1706516) IgG (H + L)-HRP, in 1X TBS-T buffer containing 1% fat free milk at a 1:3000 dilution for 2 h at RT. Immunoblots were imaged using a G:BOX Chemi XX6 system (Syngene) after adding SuperSignal West Femto reagents (Thermo Scientific, ref. 34095) directly to the PVDF membrane. Relative protein abundances were estimated with help of Fiji software (19) as follows: signal intensity of each band was calculated in the software and then normalized to the corresponding Anti-TUBULIN signal in the same running line. At least three independent biological replicates for the same experiment were done for each immunoblot.

### Co-Immunoprecipitation of proteins expressed in *Nicotiana benthamiana*

To test the protein interaction between RAPTOR1B and GIGANTEA (GI) *in planta*, the complete coding sequences of both genes were amplified using *cDNA* prepared from *A. thaliana* Col-0 plants. For *Pro35S::6xMyc-RAPTOR1B* construct, previously cloned *6xMyc-RAPTOR1B* in *pE3n* for the complementation lines in *A. thaliana* (see above) *was* subcloned via Gateway into the plant destination vector *pMDC32-HPB* (Addgene, https://www.addgene.org), which harbors a 2X CaMV 35S promoter upstream of the insertion cassette to generate *Pro35S::6XMyc-RAPTOR1B*. For *GIGANTEA, Pro35S::GI-3XHA* construct was generated by cloning first into *pE2c* and subsequently into *pMDC32-HPB* as described for *RAPTOR1B*. Both constructs were transformed into *A. tumefaciens* (strain GV3101). For *N. benthamiana* leaf infiltration, *A. tumefaciens* harboring the constructs were co-infiltrated into 6-week old plants following the protocol described in (20). 48 h after the infiltration, leaves were cut and snap frozen into liquid nitrogen and stored at -80 degrees. Subsequently, by using a mortar and pestle, the plant material was finely grinded in liquid nitrogen and 1 g of powder was resuspended into 4 ml of pre-cold extraction buffer (25 mM Tris-HCl, pH 7.6; 15 mM $MgCl_2$; 150 mM NaCl; 15 mM $pNO_2$-Phenyl$PO_4$; 60 mM B-glycerophosphate; 0.1 % NO-40 (v/v); 0.1 mM Na3$VO_4$; 1 mM NaF; 1 mM PMSF; 1 µM E64; 5 % Ethyleenglycol; 100 µM MG132; EDTA-free Ultra Complete proteases inhibitor tablets (Roche)) and mixed well by vortexing. Samples were kept on ice for 20 min and then subjected to sonication for 15 min in $dH_20$ with ice. Following, cell debris were removed by centrifugation at 4 °C the samples for 15 min. at 14000 rpm, and repeating this step 4 times more while recovering always the supernatant without disturbing the pellet. Next, for the immunoprecipitation step, the initial 4 ml suspension was divided into two 1.5 ml suspensions to be incubated separately with 50 µl of either Anti-Myc (Miltenyi Biotec, ref. 130-091-123) or Anti-HA (Miltenyi Biotec, ref. 130-091-122) microbeads for 1 h at 4 °C under continuous rotation (around 600 µl were kept apart to be used as input). Following, protein complexes were isolated using MACS columns (Milteny Biotec, ref. 130-042-701) as suggested by the manufacturer. Finally, proteins were eluted from the columns by adding pre-heated elution buffer (50 mM Tris-HCL, pH 6.8; 50 mM DTT; 1% SDS; 1 mM EDTA; 0.005% bromophenol blue; 10% glycerol) at 95°C. Co-immunoprecipitated proteins were analyzed by western blotting as described above (see “Protein extraction and immunoblot analysis”). Anti-MYC (Invitrogen, ref. 46-0603; 1:1000) and Anti-HA (Sigma, ref.H6908; 1:1000) primary antibodies were used for western blotting detection.

### Protein localization by transient expression in *Nicotiana benthamiana*

Full coding sequence of *RAPTOR1B* with stop codon was cloned into *pE3n* vector via *In-Fusion*® (Addgene, https://www.addgene.org). In the *pE3n*, 6XMyc was replaced via *In-Fusion*® cloning by *mTurquose* to generate *mTurquoise-RAPTOR1B* . For *GIGANTEA* (*GI*), full coding sequence without stop codon was cloned into *pE2C* vector via *In-Fusion*® (Addgene, https://www.addgene.org). HA tag downstream of *GI* was replaced by mNeonGreen using *In-Fusion*® to generate *GI-mNeonGreen*. Both *mTurquose-RAPTOR1B* and *GI-mNeonGreen* were separately recombined via Gateway cloning into the plant destination vector *pMDC32-HPB* to generate *Pro35S::mTurquose-RAPTOR1B* and *Pro35S::GI-mNeonGreen* (Addgene, https://www.addgene.org). Both constructs were transformed into *A. tumefaciens* for further infiltration in *N. benthamiana*. For signal detection, mTurquose was exited by a 405 nm Diode laser and signal was recovered between 455-508 nm. mNeonGreen was exited by a 488 nm Argon laser and signal was recovered between 505-560 nm. Images were acquired with the same settings for the empty vector control and respective expression constructs. Primers for cloning are listed in Table S1.

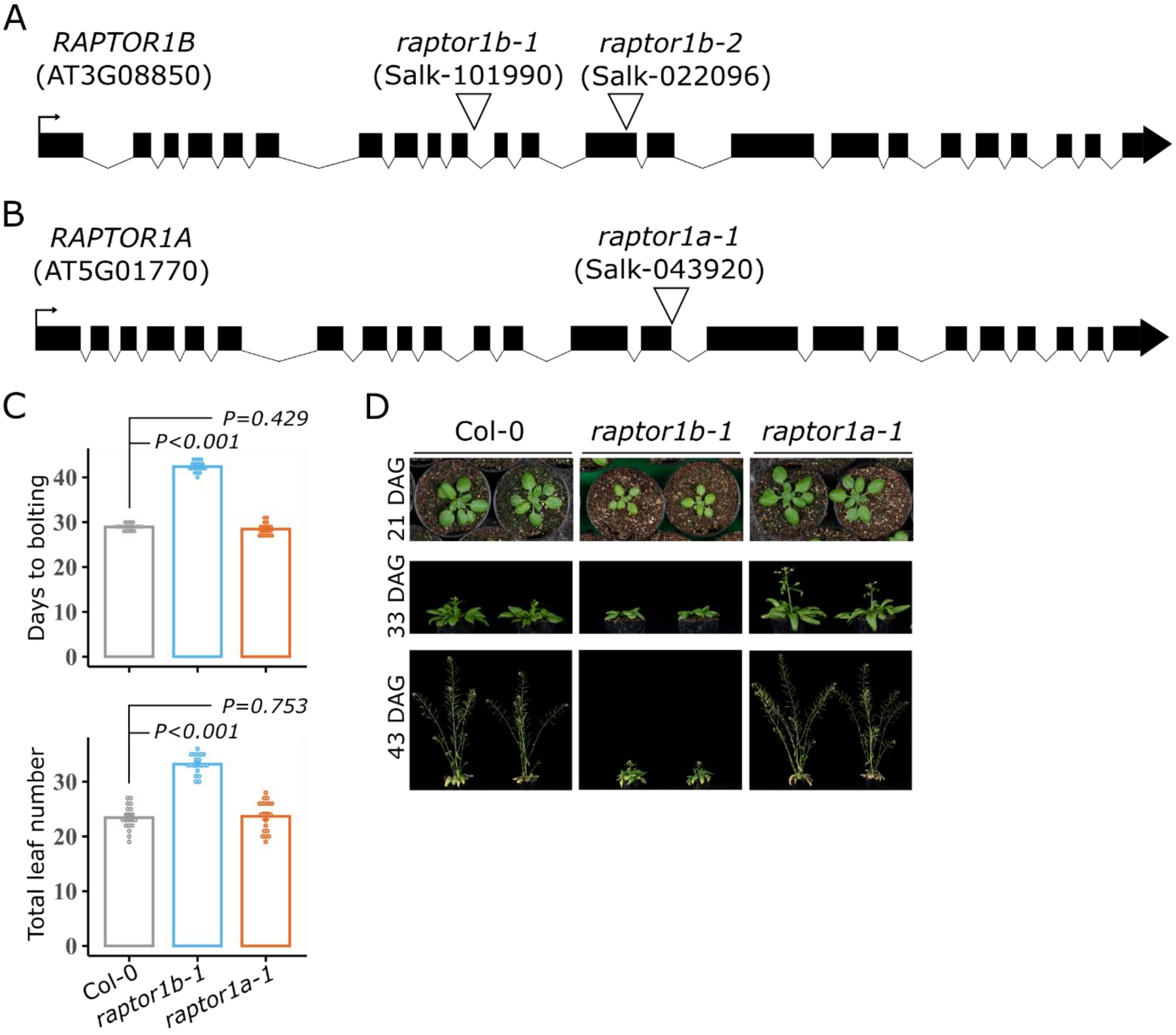


**Fig. S1. Flowering time recorded for plants with a mutant allele of *RAPTOR1A* and sketches of the T-DNA insertion lines used in this study. (A)** T-DNA insertion sites for *raptor1b-1* and *raptor1b-2* in the *RAPTOR1B* gene. **(B)** T-DNA insertion site for *raptor1a-1* in the *RAPTOR1A* gene. **(C)** Days to bolting and total leaf numbers for Col-0, *raptor1b-1* and *raptor1a-1* grown under LD conditions. Significant differences between the genotypes were determined by two-tailed Student's *t*-test (n= 20). Error bars denote SE. **(D)** Representative images of plants in (A) were taken 21, 33 and 43 days after germination (DAG).

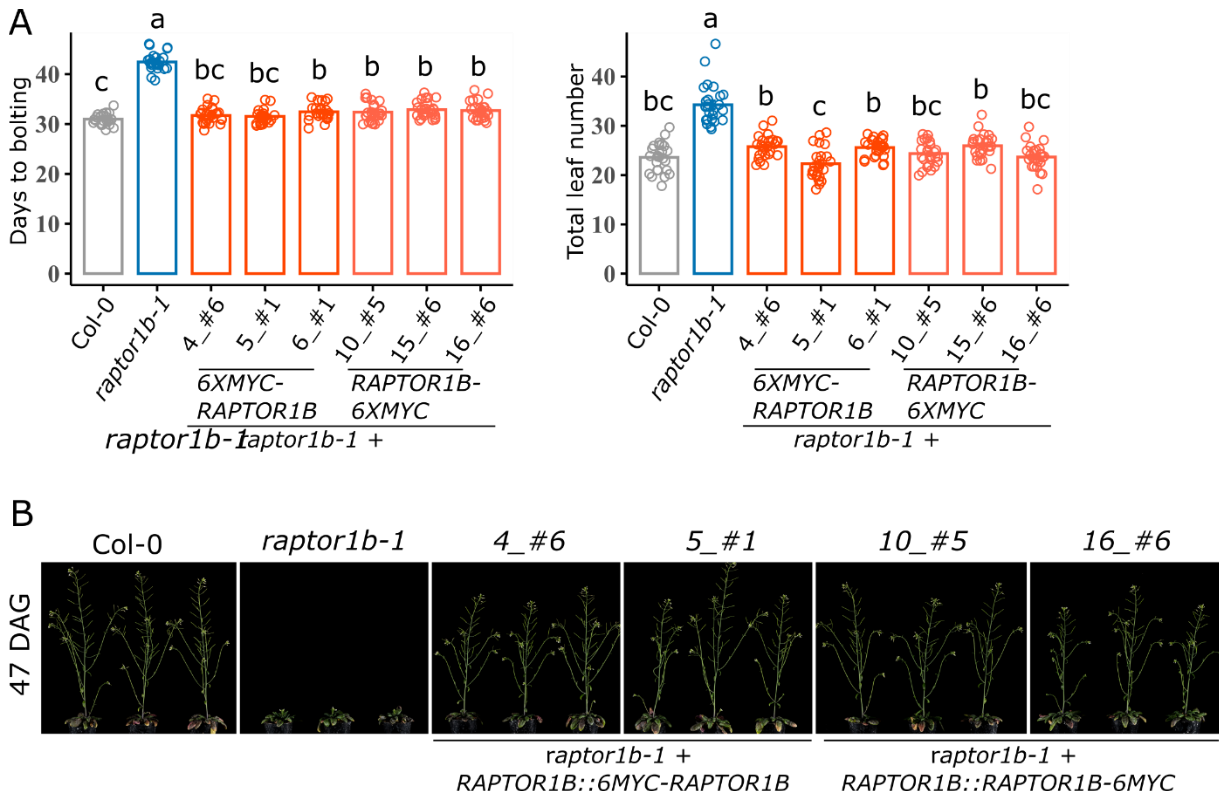


**Fig. S2. Complementation of the *raptor1b-1* mutant by stable transformatiom of *RAPTOR1B::6XMyc-RAPTOR1B* and *RAPTOR1B::RAPTOR1B-6XMyc*. (A)** Days to bolting and total leaf numbers for Col-0, *raptor1b-1* and each three independent lines complemented with *RAPTOR1B::6XMyc-RAPTOR1B* (4_#6, 5_#1 and 6_#1) and *RAPTOR1B::RAPTOR1B-6XMyc* (10_#5, 15_#6 and 16_#6) grown under LD conditions. Significant differences among genotypes were determined by one-way ANOVA ($P < 0.05$) followed by a post-hoc Tukey's test as indicated by letters (n=20). Error bars denote SE. **(B)** Representative images of Col-0, *raptor1b-1* and each two independent complemented lines used to determine flowering time in (A). Pictures were taken 47 days after germination (DAG).

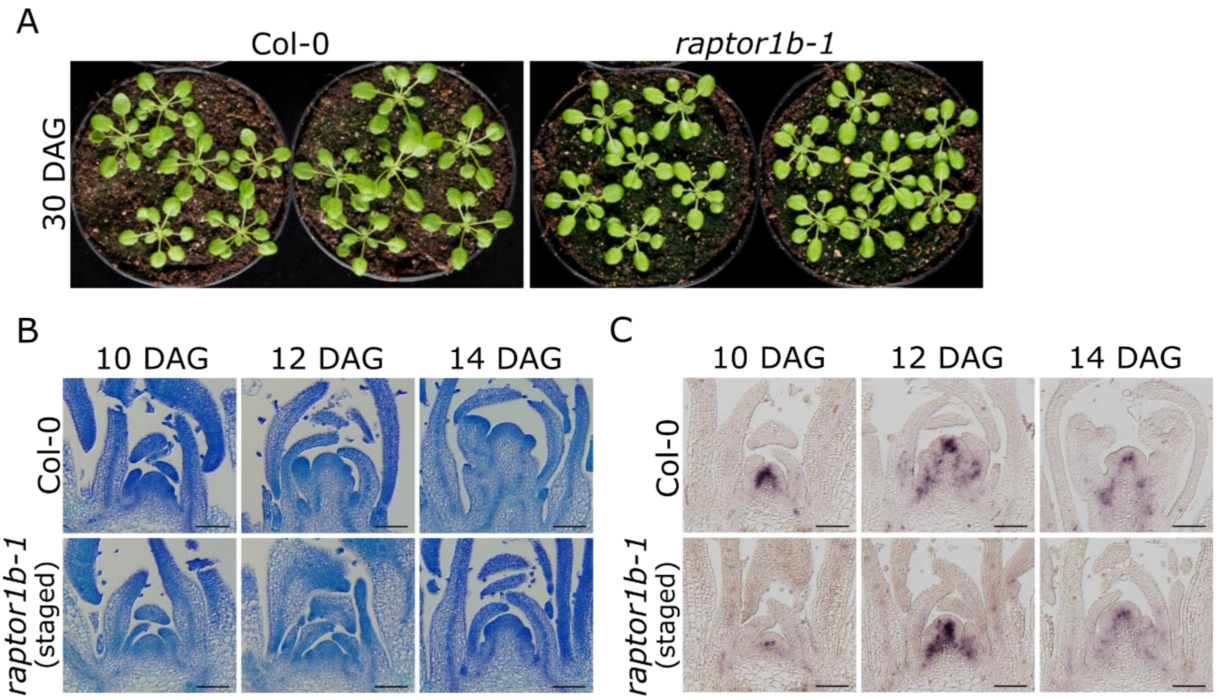


**Fig. S3. Images of Col-0 and *raptor1b-1* plants grown under SD and longitudinal sections of apices for the same genotypes grown under LD. (A)** Representative pictures of wild-type (Col-0) and *raptor1b-1* plants grown in short day (SD) conditions for 30 days. This plant material was used for the gene and protein expression analysis depicted in Figure 2A, B and C, corresponding to the 0DAS harvesting point before the photoperiod shift. **(B)** Longitudinal sections through Col-0 and *raptor1b-1* apices stained with Toluidine blue, for which plants were grown under long photoperiod and apices were harvested at 10, 12, and 14 days after germination (DAG). Given that *raptor1b* mutants display late germination (21), *raptor1b-1* seeds were brought to the growth chamber two days in advance (staged) compared to Col-0 seeds, minimizing the effect of late germination at the time of floral transition. Apices were harvested 1 h before dusk. DAG: days after germination. Scale bar: 100 µm. **(C)** RNA *in situ* hybridization using a specific probe for *SOC1* at 10, 12, and 14 DAG. Plants were grown and harvested as described in (B). Bar: 100 µm.

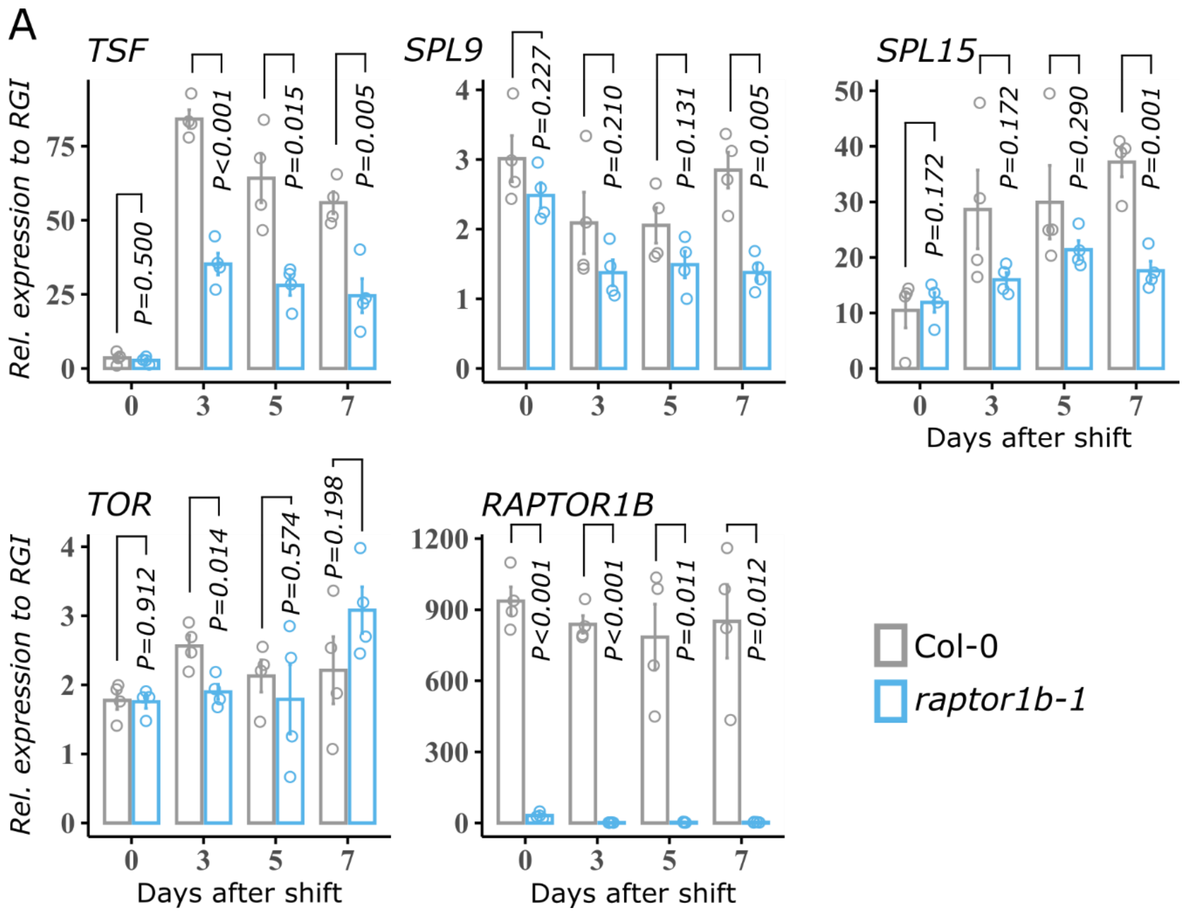


**Fig. S4. RAPTOR1B promotes the expression of flowering genes. Supplementary information of Figure 2. (A)** Gene expression analysis of *TSF*, *SPL9, SPL15, TOR and RAPTOR1B* in Col-0 and *raptor1b-1* plants by RT-qPCR. Growth conditions, plant material, harvesting time points and data analysis are described in Fig. 2A. Significant differences between the genotypes for each day were determined by two-tailed Student's *t*-test (n= 4). Error bars denote SE.

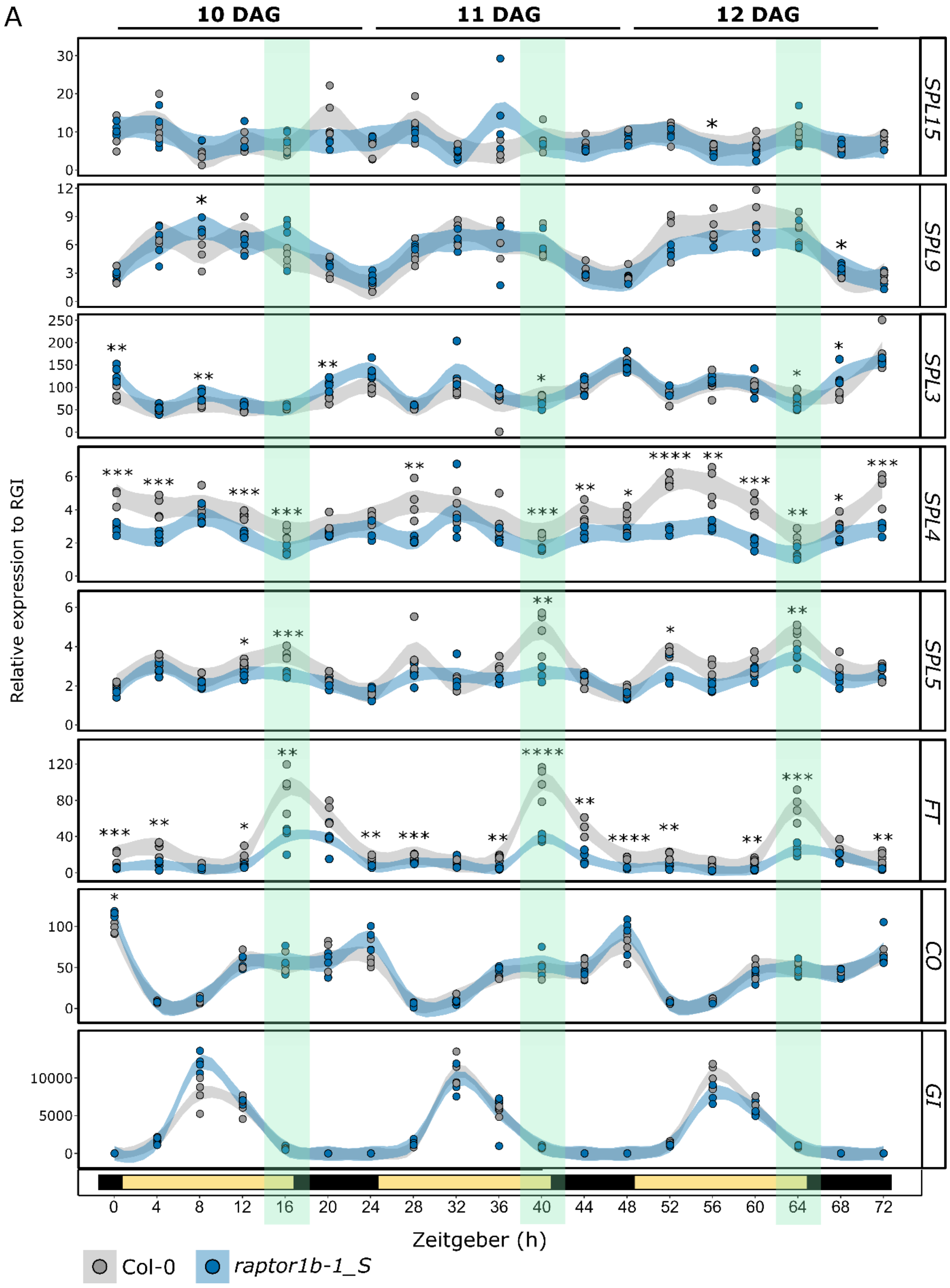


**Fig. S5. Diel expression analysis of *FT*, *CO*, *GI,* SPL3, SPL4, SPL5, SPL9 and SPL15 in Col-0 and *raptor1b-1* under LD conditions. (A)** Col-0 and *raptor1b-1* plants were grown from the beginning under LD photoperiod. *raptor1b-1* seeds were brought into the growth chamber two days in advance after stratification to compensate for the late germination (1), and therefore, it is referred in the plot as *raptor1b-1*_S (S, staged). Whole rosettes were harvested every 4 h for three consecutive days at 10, 11 and 12 DAG, which is the time when the floral transition occurs (see *SI Appendix*, Fig. S3B and C).

Four biological replicates per time point and genotype were done by harvesting a pool of ~5 rosettes in each case. Relative gene expression was calculated as explained in Fig. 2A. Significant differences between the genotypes for each time point were determined by two-tailed Student's *t-test* (n= 4, *$P<0.05$, **$P<0.01$ , ***$P<0.001$, ****$P<0.0001$). Error bars denote SE. The shaded green-field area along the plots highlights the *Zeitgeber* time 16 h (dusk) for each day.

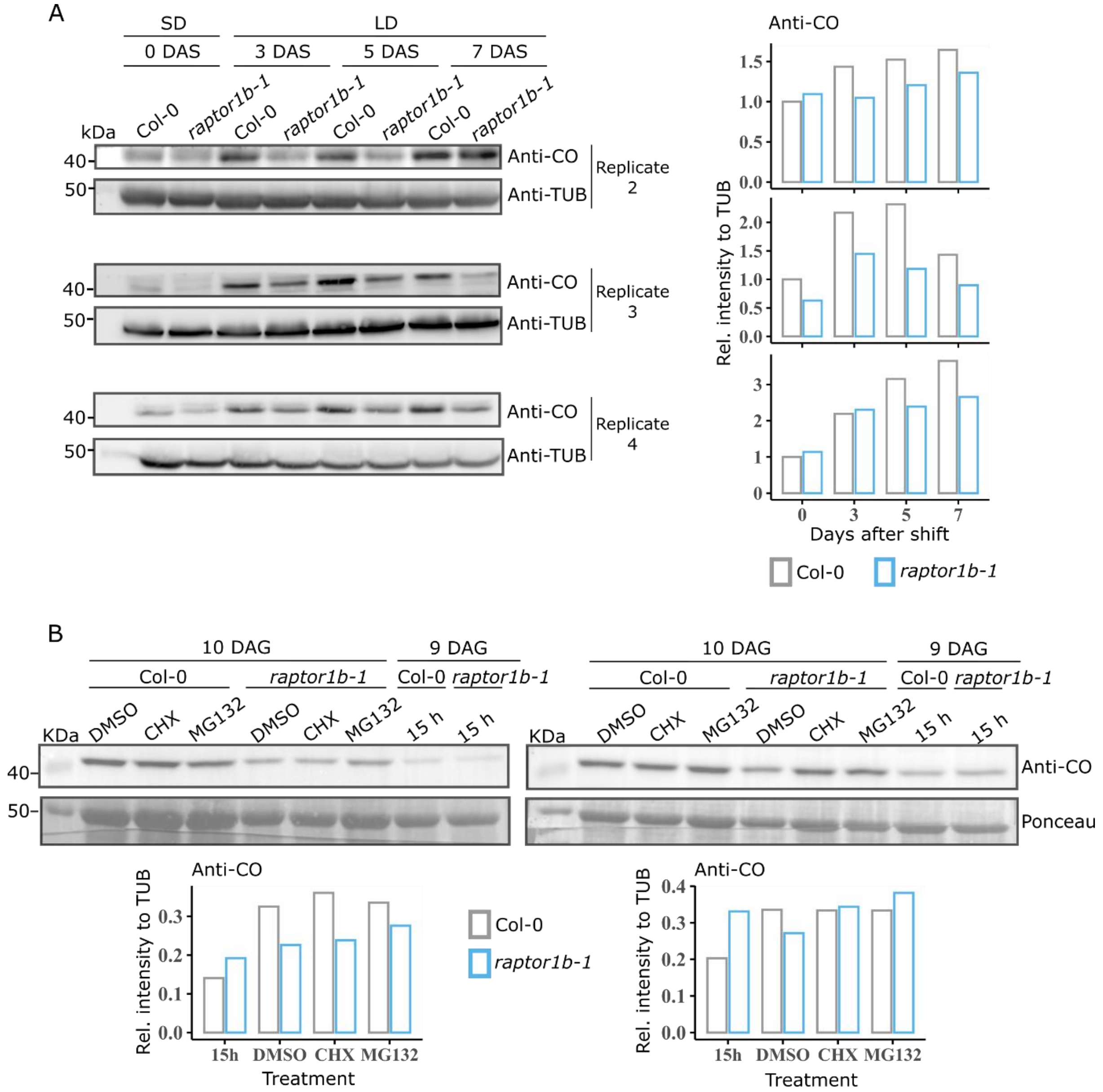


**Fig. S6. RAPTOR1B contributes to CONSTANS protein stability under LD at the time of the floral transition. Supplementary information of Fig. 3**. **(A)** Additional replicates of the western blot analysis to determine CONSTANS protein levels described in main Fig. 3A. On the right panel, relative CONSTANS protein levels correspond to each of the replicates on the left side and the relative intensities were calculated as described in Fig. 3B. **(B)** Additional replicates of the immunoblot analysis of CONSTANS protein levels in Col-0 and *raptor1b-1* plants upon treatment with Cycloheximide (CHX) or MG132 as described in main Fig. 3C. Below each blot, a plot depicts the relative CONSTANS protein levels calculated as described in Fig. 3D.

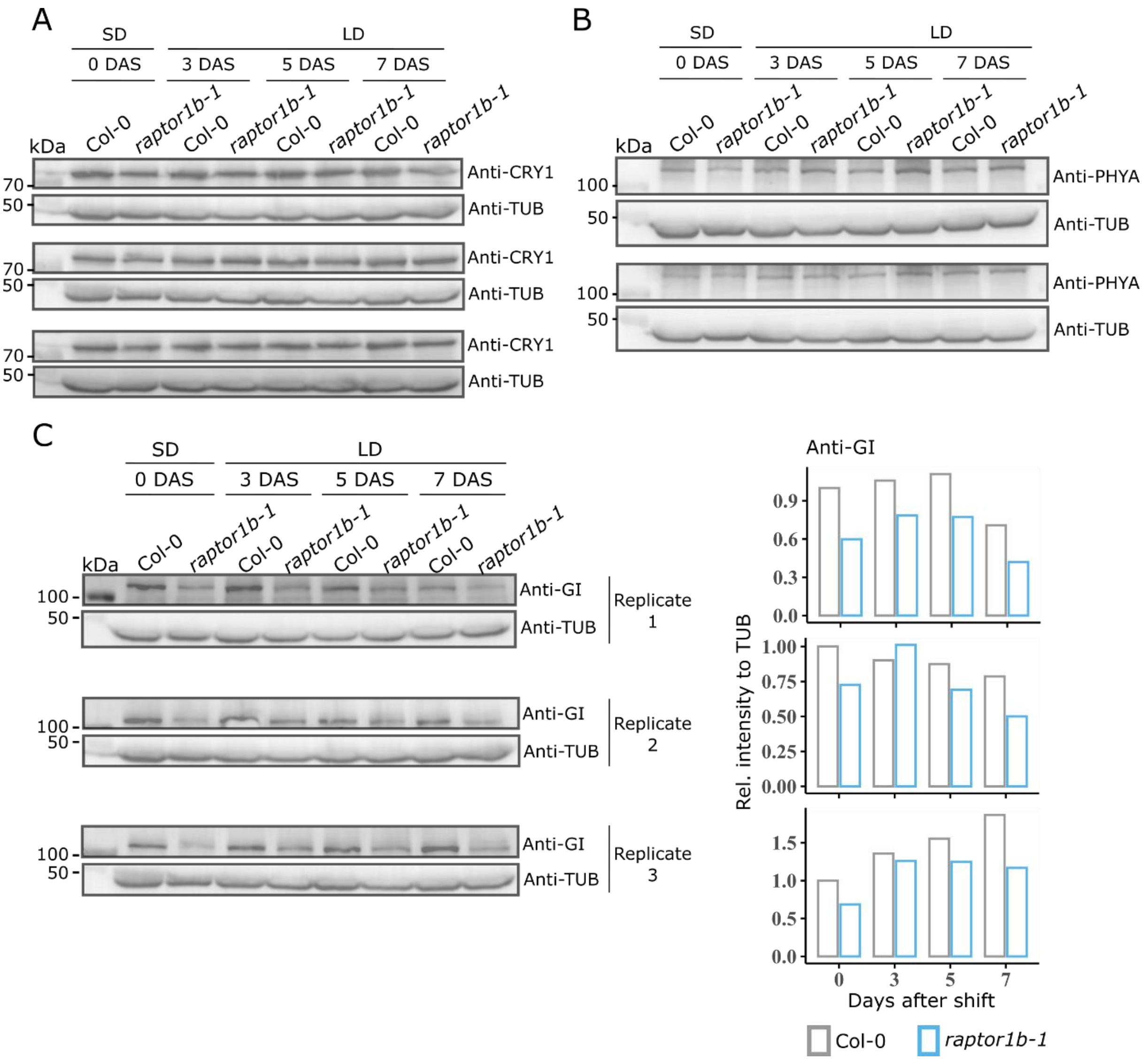


**Fig. S7. Western blot analysis to determine protein levels of CRYPTOCHROME 1 (CRY1), PHYTOCHROME A (PHYA) and GIGANTEA (GI) in Col-0 and *raptor1b-1* during a SD to LD photoperiod shift. (A)** Western blot for CRY1. Grow conditions and plant material are the same as described in Fig. 2A. Proteins were extracted and endogenous CRY1 and TUBULIN proteins were immuno-detected using Anti-CRY1 and Anti-TUB, respectively. Three biological replicates are depicted. **(B)** Western blot for PHYA. Grow conditions and plant material are the same as described in Fig. 2A. Proteins were extracted and endogenous PHYA and TUBULIN proteins were immuno-detected using Anti-PHYA and Anti-TUB, respectively. Two biological replicates are depicted. **(C)** Additional replicates of the western blot analysis to determine GIGANTEA (GI) protein levels described in main Fig. 4A. On the right panel, relative GI levels correspond to each of the replicates on the left side and the relative intensities were calculated as described in Fig. 4B.

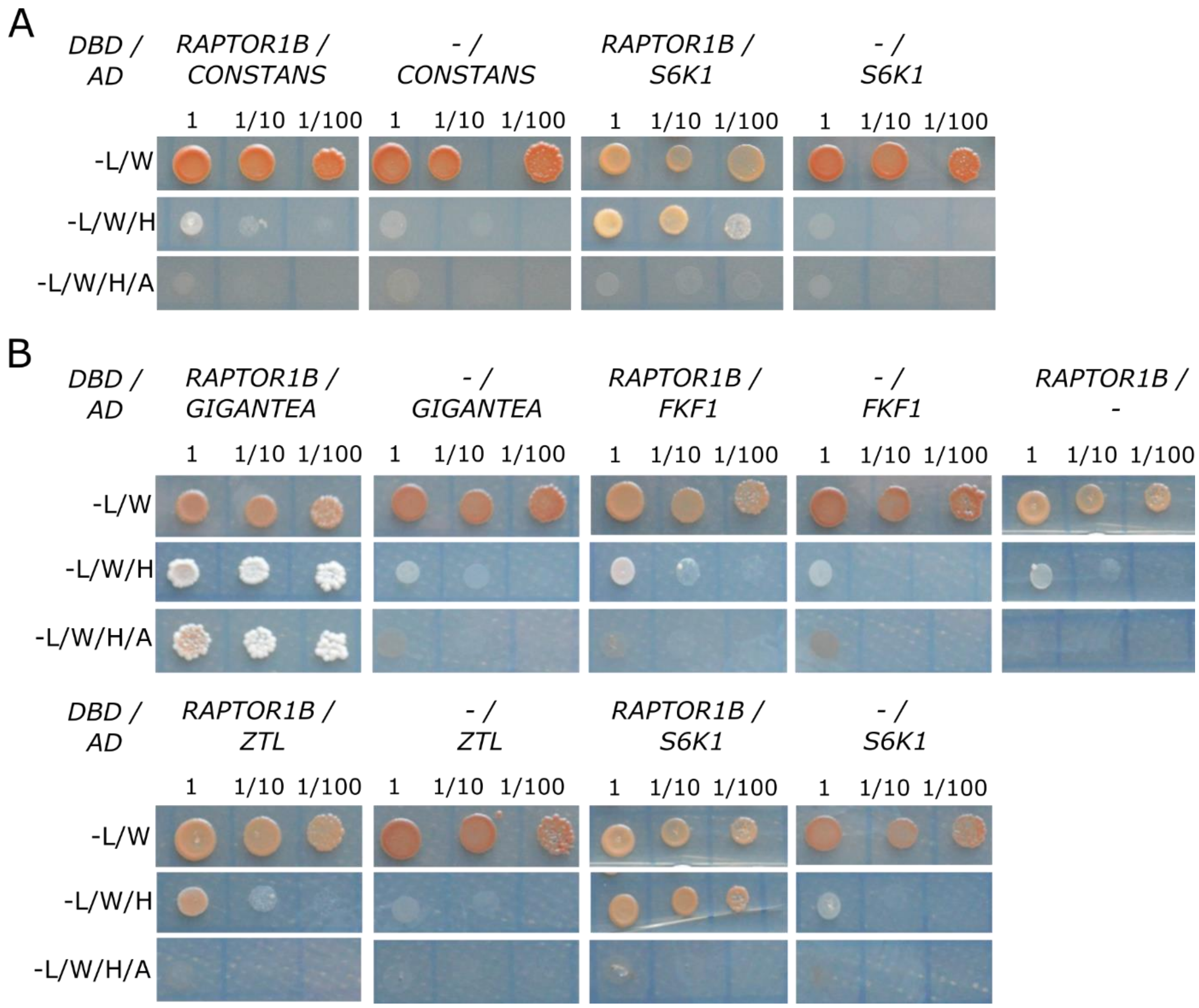


**Fig. S8. Yeast 2 hybrid assays to test additional interactions of RAPTOR1B, including the positive control with 40S Ribosomal protein S6 Kinase 1 (S6K1). (A)** Full coding sequence of *RAPTOR1B* was cloned into *pDEST™32* (harboring the DNA binding domain, DBD). Full coding sequence of *CO* and a cDNA fragment from amino acid 9 to 293 in case of *S6K1*, were cloned into *pDEST™22* (containing the activation domain, AD). *pDEST™32* and *pDEST™22* constructs were co-transformed into yeast following the protocol in (14). **(B)** *RAPTOR1B* and *S6K1* were cloned as in (A). Full coding sequences of *GIGANTEA*, *FKF1*, and *ZTL* were cloned into *pDEST™22*. Constructs were co-transformed as in (A). In (A) and (B), as controls, respective empty *pDEST™32* lacking *RAPTOR1B* or *pDEST™22* lacking the other proteins were simultaneously co-transformed and spotted in the same plate. The different combinations of plasmids were spotted on synthetic double (- L/W), triple (- L/W/H) or quadruple (- L/W/H/A) dropout medium. L (Leucine), W (Tryptophan), H (Histidine), A (Adenine).

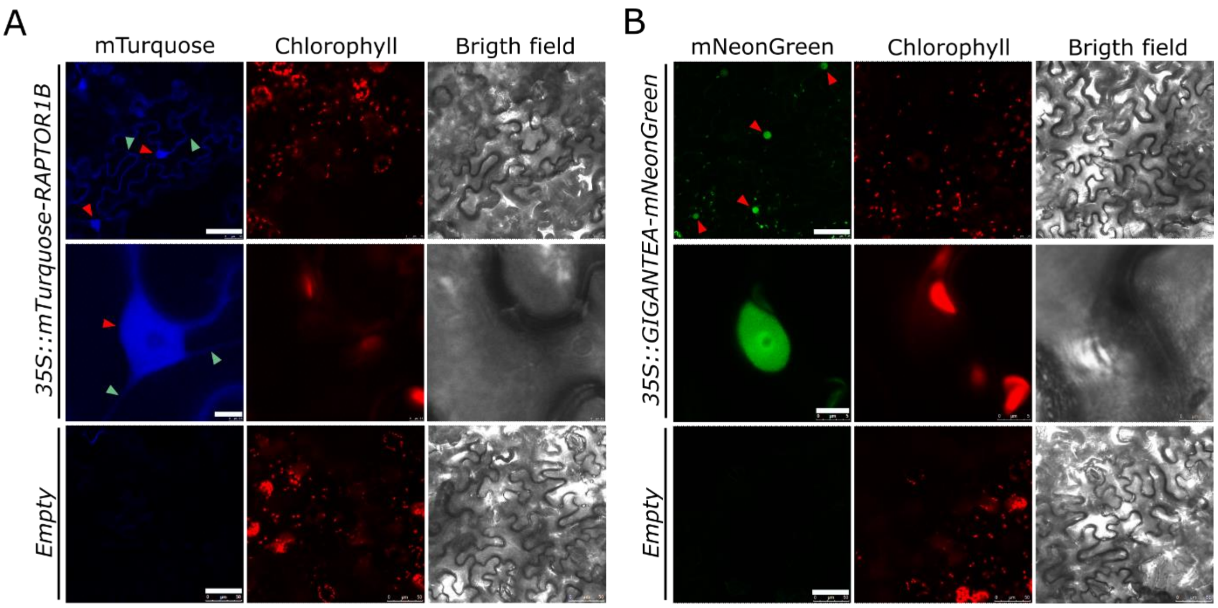


**Fig. S9. Subcellular localization of RAPTOR1B and GIGANTEA. (A)** Transient expression of *Pro35S::mTurquoise-RAPTOR1B* in *N. benthamiana* leaves. Upper panel depicts mTurquoise signal in nuclei (red triangles) and in cytoplasm (green triangles). Middle panel displays a closer view of a selected nucleus (red triangle) and cytoplasmic string are also visible (green triangle). Bottom panel corresponds to infiltration with the empty vector backbone (*pMDC32-HPB*). All images were acquired using the same settings for detecting mTurquoise signal. **(B)** Transient expression of *Pro35S::GI-mNeonGreen* in *N. benthamiana* leaves. Upper panel depicts mNeonGreen signal in nuclei (red triangles). Middle panel displays a closer view of a selected nucleus. Bottom panel corresponds to infiltration with the empty vector backbone (*pMDC32-HPB*). All images were acquired using the same settings for detecting mNeonGreen signal. Scale bars for (A) and (B), upper panels: 50 µm, middle panels: 5 µm, bottom panels: 50 µm.

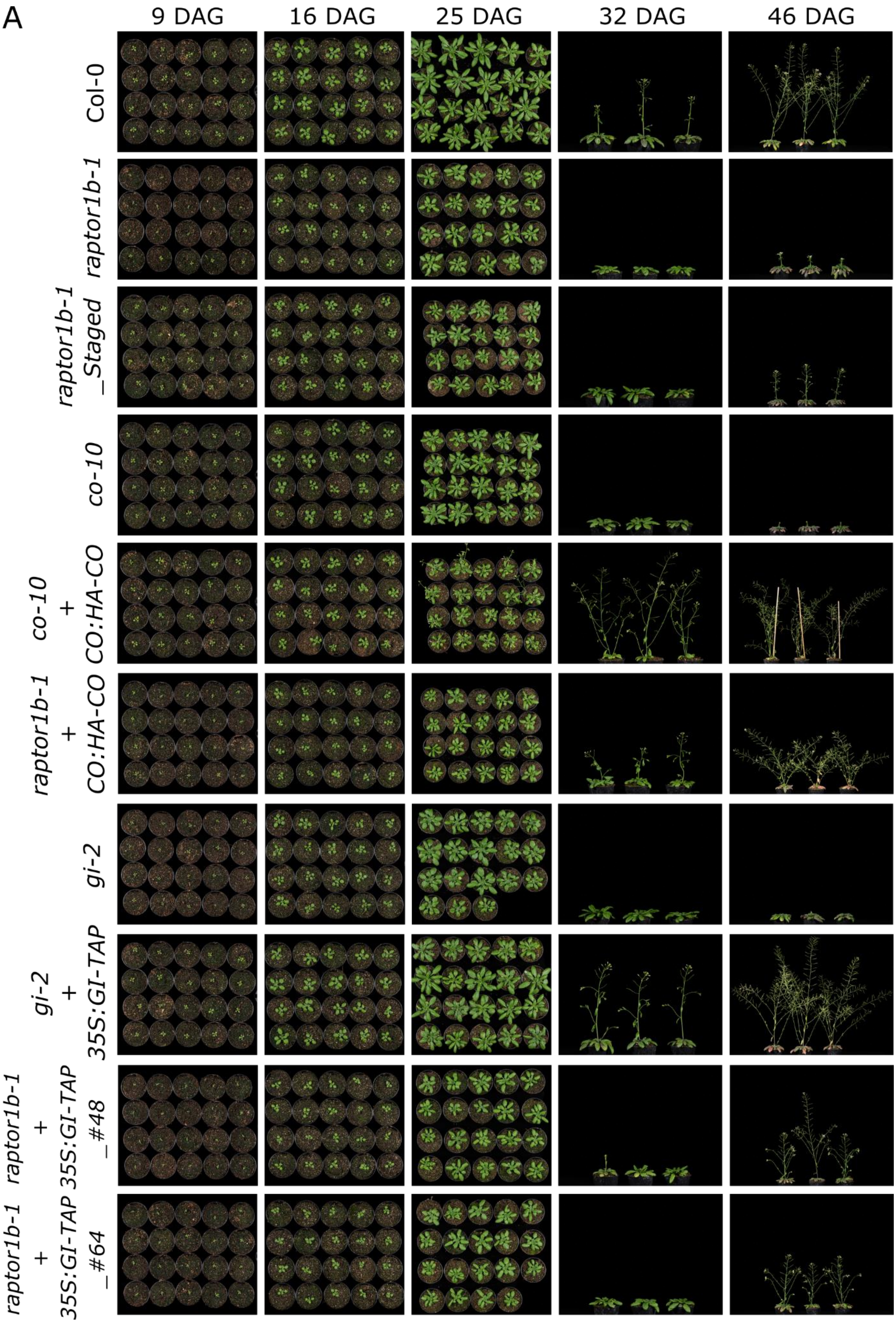


**Fig. S10. (A)** Images corresponding to the flowering time experiment in Fig. 5A. Images were taking 9, 16, 25, 32 and 46 after germination (DAG).

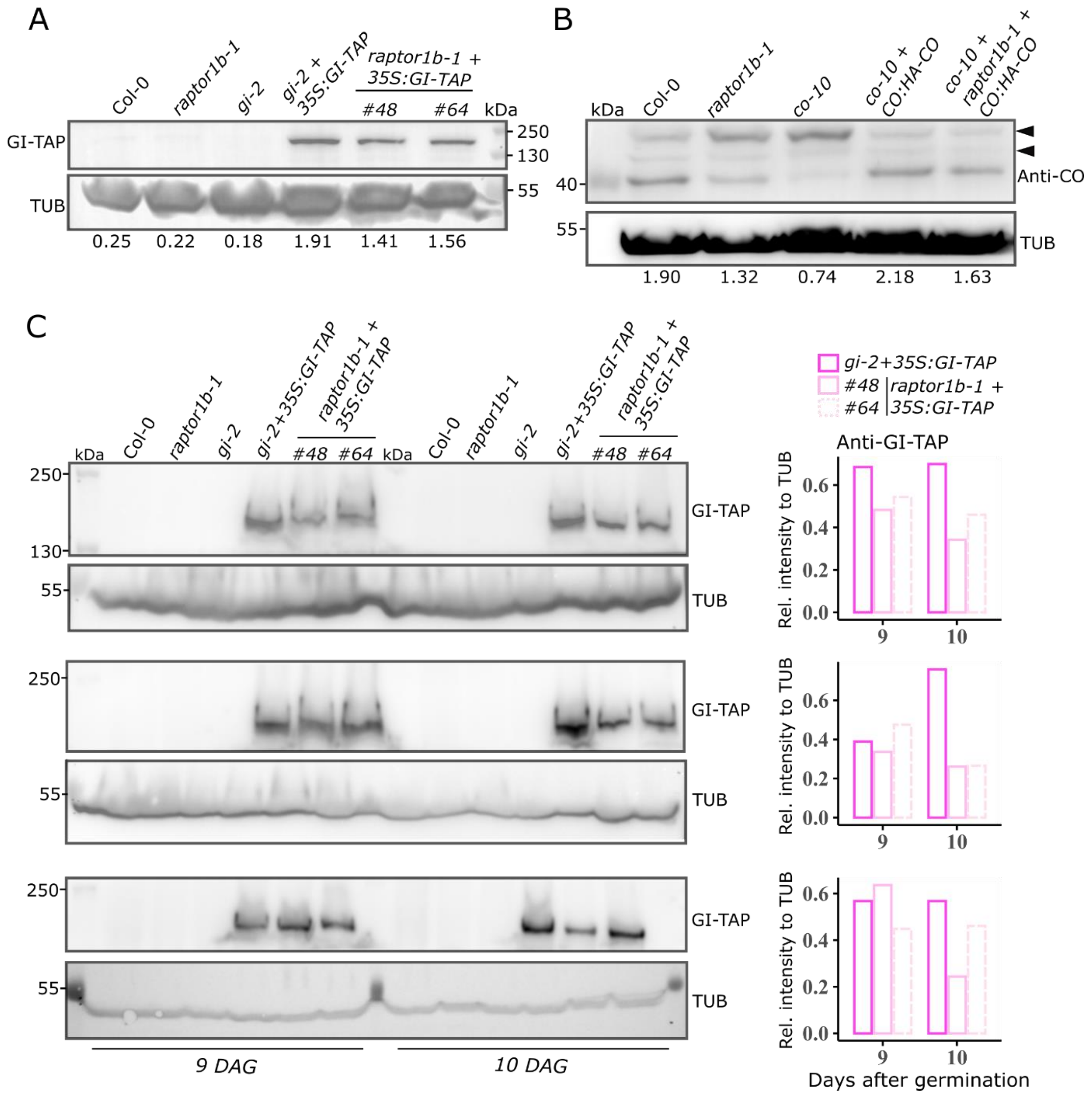


**Fig. S11. Western blot analyses for GIGANTEA and CONSTANS at bolting and during the floral transition. (A)** Western blot analysis for GI-TAP (~150 kDa). Plant material corresponds to flowering time experiment of main Fig. 5A. At bolting, for each genotype, a pool of three rosettes were collected 15 h after the onset of light. Proteins were extracted and GI-TAP and TUBULIN were immuno-detected using Anti-GI and Anti-TUB, respectively. Anti-GI signal was normalized to the respective Anti-TUB signal to determine the relative intensity for each genotype and the respective numbers are depicted below the blots. **(B)** Western blot analysis for CONSTANS (CO) (endogenous CO ~42 kDa; HA-CO ~43 kDa). Plant material corresponds to main Fig. 5A and protein extraction was performed as described in (A). CO/HA-CO and TUBULIN were immuno-detected using Anti-CO and Anti-TUB, respectively. Anti-CO signal was normalized to the respective Anti-TUB signal to determine the relative intensity for each genotype and the respective numbers are depicted below the blots. Dark triangles depict unspecific bands. Note: HA-tag adds 1.1 kDa to endogenous CO. **(C)** Additional replicates of the western blot analysis described in main Fig. 5B and C. On the right panel, relative GI levels correspond to each of the replicates on the left side and the relative intensities were calculated as described in Fig. 5C.

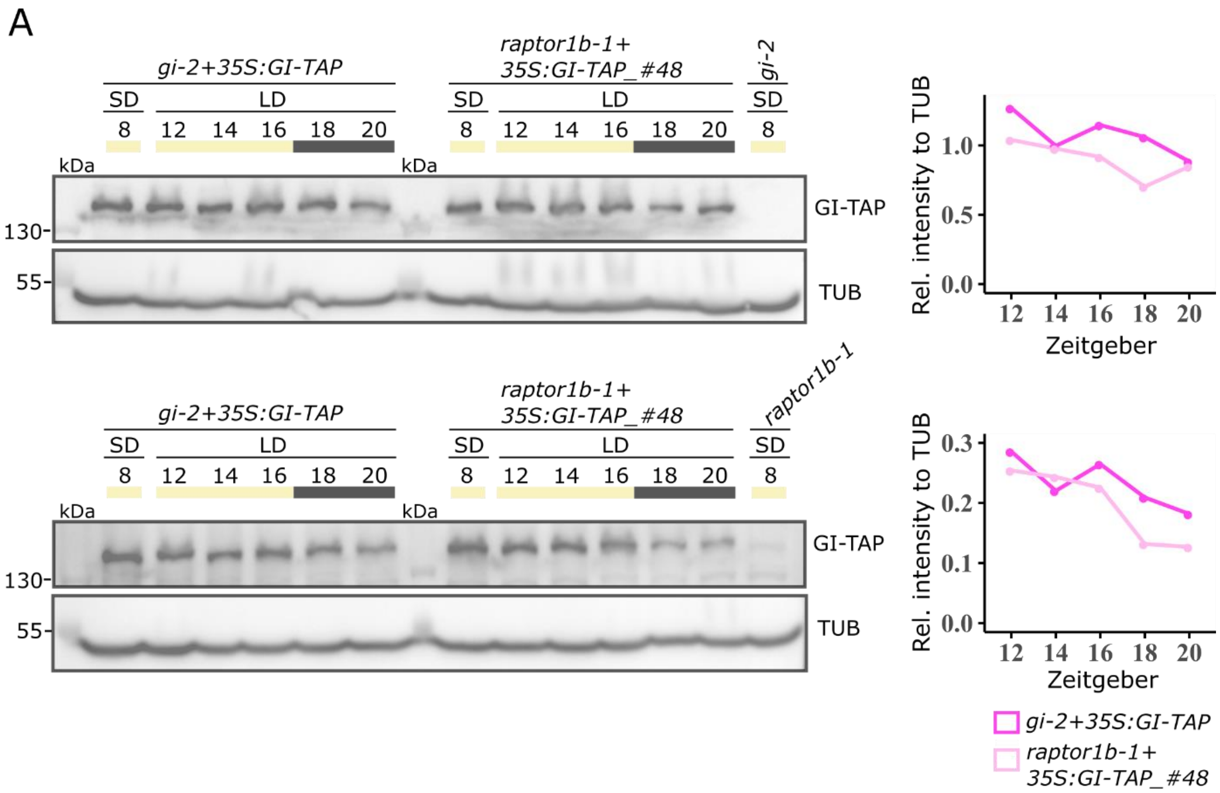


**Fig. S12. (A)** Additional replicates of main Fig. 5D and E. For these replicates, instead of Col-0, proteins were extracted from *gi-2* or *raptor1b-1* mutants to be used as controls, respectively. The two plots on the right correspond to the relative GI-TAP levels calculated from the blots on the left panel as described in main Fig. 5E.

**Table S1. Primer list**

| AGI /Gene bank | Primer Name | Sequence | Purpose |
|---|---|---|---|
| | BP | attttgccgatttcggaac | Genotyping SALK lines |
| | LB-1 | tagcatctgaatttcataaccaatctcgatacac | Genotyping SAIL lines |
| AT3G08850 | RAPTOR B LP_3 | agcagtgacggatacaattgc | Genotyping *raptor1b-1* (SALK_101990) |
| AT3G08850 | RAPTOR B RP_3 | gttgtttgcagaagcttcagc | Genotyping *raptor1b-1* (SALK_101990) |
| AT3G08850 | RAPTOR B LP_4 | caatatgaagctgcggctaac | Genotyping *raptor1b-2* (SALK_022096) |
| AT3G08850 | RAPTOR B RP_4 | catcggatcaagttgcttacc | Genotyping *raptor1b-2* (SALK_022096) |
| AT5G01770 | ra1_LP | aagaggatgagcgaactaggc | Genotyping *raptor1a-1* (SALK_043920) |
| AT5G01770 | ra1_RP | tttgtcttggaaagttgtggg | Genotyping *raptor1a-1* (SALK_043920) |
| At5g15840 | co_LP | aagctgttgtgacacatgctg | Genotyping *co-10* (SAIL_24_H04) |
| At5g15840 | co_RP | ccccttctttcagataccagc | Genotyping *co-10* (SAIL_24_H04) |
| AT1G22770 | gi-2_Fw_seq500bp | ttcatgtcttattga | Genotyping/sequencing for gi-2 deletion |
| AT1G22770 | gi-2_Rv_seq500bp | aaataaaagaaggag | Genotyping/sequencing for gi-2 deletion |
| AT3G08850 | RAPTORB_Forw_pE3n | cggaagcccgggatccatggcattaggagacttaatggtg | Cloning ProRAPTOR1B::6XMyc-RAPTOR1B, Co-Immunoprecipitation and subcellular localization |
| AT3G08850 | RAPTORB_Rev_pE3n | gctacttatgcggccgctcatcttgcttgcgagttgtc | Cloning ProRAPTOR1B::6XMyc-RAPTOR1B, Co-Immunoprecipitation and subcellular localization |
| AT3G08850 | RAPTORB_Forw_pE3c | tcagtcgactggatccatggcattaggagacttaatggtg | Cloning ProRAPTOR1B::RAPTOR1B-6XMyc |
| AT3G08850 | RAPTORB_Rev_pE3c | atagcttgtgcggccgctcttgcttgcgagttgtc | Cloning ProRAPTOR1B::RAPTOR1B-6XMyc |
| AT3G08850 | promRAP fwB | accaattcaggtcgacgatgcgagattcccaagtgagagag | Cloning ProRAPTOR1B (Promoter) |
| AT3G08850 | promRAP revB | aatcgataccgtcgacgaaatcacgcaaatcaaatcctcagcc | Cloning ProRAPTOR1B (Promoter) |
| AT3G08850 | RAPTORB_cDNA_Forw | ggggacaagtttgtacaaaaaagcaggcttttggatggcattaggagacttaatg | Cloning for Yeast two hybrid |
| AT3G08850 | RAPTORB_cDNA_Rev | ggggaccactttgtacaagaaagctgggtcgactcatcttgcttgcgagttgt | Cloning for Yeast two hybrid |
| At5g15840 | CO_cDNA_Forw | ggggacaagtttgtacaaaaaagcaggctttatgttgaaacaagagagtaacgaca | Cloning for Yeast two hybrid |
| At5g15840 | CO_cDNA_Rev | ggggaccactttgtacaagaaagctgggtatcagaatgaaggaacaatcccata | Cloning for Yeast two hybrid |
| AT1G22770 | GI_cDNA_Forw | ggggacaagtttgtacaaaaaagcaggctggatggctagttcatcttcatctga | Cloning for Yeast two hybrid |
| AT1G22770 | GI_cDNA_Rev | ggggaccactttgtacaagaaagctgggtgttattgggacaaggatatagtacagcc | Cloning for Yeast two hybrid |
| AT1G68050 | FKF1_cDNA_Forw | ggggacaagtttgtacaaaaaagcaggctttatggcgagagaacatgcga | Cloning for Yeast two hybrid |

| | | | |
|---|---|---|---|
| AT1G68050 | FKF1_cDNA_Rev | ggggaccactttgtacaagaaagctgggttttacagatccgagtcttgccg | Cloning for Yeast two hybrid |
| AT5G57360 | ZTL_cDNA_Forw | ggggacaagtttgtacaaaaaagcaggctttatggagtgggacagtggttc | Cloning for Yeast two hybrid |
| AT5G57360 | ZTL_cDNA_Rev | ggggaccactttgtacaagaaagctgggttctaatgaggaagaaagaagaagaagga | Cloning for Yeast two hybrid |
| AT3G08730 | S6K1_Forw_Peptide | ggggacaagtttgtacaaaaaagcaggctttcccaacaaaatccagaaacagc | Cloning for Yeast two hybrid |
| AT3G08730 | S6K1_Rev_Peptide | ggggaccactttgtacaagaaagctgggtctggctatccgcacatggagtttgatc | Cloning for Yeast two hybrid |
| AT1G22770 | GI_HA_C_For | tcagtcgactggatccatggctagttcatcttcatctgag | Cloning for Co-Immunoprecipitation and subcellular localization |
| AT1G22770 | GI_HA_C_Rev | cctccgcttgcggccgcttgggacaaggatatagtacagccg | Cloning for Co-Immunoprecipitation and subcellular localization |
| At5g20700 | gDNA FLZ14 N fw | ggccgctggggccatgcttactaaaagaacccatc | Cloning for Co-Immunoprecipitation (Control) |
| At5g20700 | attB1 mNEON C FLZ14 fw | ggggacaagtttgtacaaaaaagcaggcttaatgcttactaaaagaacccatcccatga | Cloning for Co-Immunoprecipitation (Control) |
| UEC50308.1 | mNeonGreen tag F | ggggacaagtttgtacaaaaaagcaggcttaatggtgagcaagggagagga | Cloning for Co-Immunoprecipitation (Control) |
| UEC50308.1 | mNeonGreen tag R | ttttagtaagcatggccccagcggccgca | Cloning for Co-Immunoprecipitation (Control) |
| QBQ65841.1 | Tourq_For_N_2_RAPTOR | accaattcaggtcgacatggtgagcaagggcgagga | Cloning for subcellular localization |
| QBQ65841.1 | Tourq_Rev_N_2_RAPTOR | tgccatggatcccggggaggccccagcggccgca | Cloning for subcellular localization |
| UEC50308.1 | mNeon_For_C_2_GI | ttgtcccaagcggccgcaggctcgggaggtggagg | Cloning for subcellular localization |
| UEC50308.1 | mNeon_Rev_C_2_GI | gaaagctgggtctagatttgtatagctcgtccattcccatc | Cloning for subcellular localization |
| At1G65480 | FT_F | tggaacaacctttggcaatgag | qRT-PCR |
| At1G65480 | FT_R | cgacacgatgaattcctgcag | qRT-PCR |
| AT4G20370 | TSF_F | ctcgggaattcatcgtattg | qRT-PCR |
| AT4G20370 | TSF_R | ccctctggcagttgaagtaa | qRT-PCR |
| AT2G33810 | SPL3_F | gagtttgtcaggtcgagagttgtacc | qRT-PCR |
| AT2G33810 | SPL3_R | gcagactttgtgtcgtttgtggt | qRT-PCR |
| AT1G53160 | SPL4_F | aatggtcaggtggtgatgcag | qRT-PCR |
| AT1G53160 | SPL4_R | gcataggaagtgtcatctctaccctt | qRT-PCR |
| AT3G15270 | SPL5_F | cagcaggtttcatgagctaccag | qRT-PCR |
| AT3G15270 | SPL5_R | caaaactgtcaccagagatcttcctc | qRT-PCR |
| AT2G42200 | SPL9_F | cttcgctttacgaaaatggtgatg | qRT-PCR |
| AT2G42200 | SPL9_R | actggccgcctcatcactct | qRT-PCR |
| AT3G57920 | SPL15_F | catctctttacggaaaccccaatg | qRT-PCR |
| AT3G57920 | SPL15_R | gccgctgcatcactgatctt | qRT-PCR |
| AT1G22770 | GI_F | agcagtggtcgacggtttatc | qRT-PCR |
| AT1G22770 | GI_R | atgggtatggagctttggttc | qRT-PCR |
| At5g15840 | CO_F | aacagcttcacacccaagaacg | qRT-PCR |
| At5g15840 | CO_R | ggtcaggttgttgctctactg | qRT-PCR |
| AT3G08850 | RAPTOR1B_F | tcaatccagggtcacaagcc | qRT-PCR |
| AT3G08850 | RAPTOR1B_R | gatgcactcaccacctttgc | qRT-PCR |
| AT1G50030 | TOR_F | catctgcgcgtctggaaatg | qRT-PCR |
| AT1G50030 | TOR_R | cctcgtcgtactttgccctt | qRT-PCR |
| AT1G13320 | PDF2_F | taacgtggccaaaatgatgc | qRT-PCR |
| AT1G13320 | PDF2_R | gttctccacaaccgcttggt | qRT-PCR |
| AT4G34270 | TIP41_F | gtgaaaactgttggagagaagcaa | qRT-PCR |

| | | | |
|---|---|---|---|
| AT4G34270 | TIP41_R | tcaactggatacccctttcgca | qRT-PCR |
| AT4G27960 | UBC9_F | tcacaatttccaaggtgctgc | qRT-PCR |
| AT4G27960 | UBC9_R | tcatctgggtttggatccgt | qRT-PCR |
| AT4G26410 | RHIP_F | gagctgaagtggcttccatgac | qRT-PCR |
| AT4G26410 | RHIP_R | ggtccgacatacccatgatcc | qRT-PCR |
